# The subclonal footprint of pervasive early dissemination in pancreatic cancer

**DOI:** 10.1101/2025.11.27.690823

**Authors:** Yuanchang Fang, Michelle Chan-Seng-Yue, Karen Ng, Amy Zhang, Tuan Hoang, Gun Ho Jang, Sabiq Chaudhary, Catia Gaspar, Eugenia Flores-Figueroa, Daniela Bevacqua, Stephanie Ramotar, Ayelet Borgida, Shawn Hutchinson, Anna Dodd, Ayah Elqaderi, Maryam Monajemzadeh, Yifan Wang, Stanley W. K. Ng, Spring Holter, Barbara T. Grünwald, Julie M. Wilson, Robert C. Grant, Erica S. Tsang, George Zogopoulos, Masoom A. Haider, Grainne M. O’Kane, Jennifer J. Knox, Steven Gallinger, Faiyaz Notta

**Affiliations:** Princess Margaret Cancer Centre, Toronto, Ontario, Canada; PanCuRx Translational Research Initiative, Ontario Institute for Cancer Research, Toronto, Ontario, Canada; Department of Medical Biophysics, University of Toronto, Toronto, Ontario, Canada; Scarborough Health Network, Toronto, Ontario, Canada; Wallace McCain Centre for Pancreatic Cancer, Princess Margaret Cancer Centre, University Health Network, Toronto, Ontario, Canada; Toronto General Hospital, Toronto, Ontario, Canada; Department of Basic Medical Sciences, Faculty of Medicine, Ibn Sina University for Medical Sciences, Al-Qastal, Jordan; Department of Pathology and Molecular Medicine, Faculty of Health Sciences, McMaster University, Hamilton, Ontario, Canada; The Research Institute of the McGill University Health Centre, Montreal, QC, Canada; The Rosalind & Morris Goodman Cancer Institute, Montreal, QC, Canada; Department of Biological Chemistry, School of Medicine, University of California, Irvine (UCI), Irvine, California, USA; Department of Urology, University Hospital Essen, Essen, Germany; West German Cancer Center, Essen, Germany; Institute of Medical Science, University of Toronto, Toronto, Ontario, Canada; Institute of Clinical Evaluative Sciences, University of Toronto, Toronto, Ontario, Canada; Joint Department of Medical Imaging, Princess Margaret Hospital, Sinai Health System, University of Toronto, Toronto, ON, Canada; Lunenfeld-Tanenbaum Research Institute, Mount Sinai Hospital, Toronto, ON, Canada; St. Vincent’s University Hospital Dublin, School of Medicine University College Dublin, Dublin, Ireland; Department of Surgery, University of Toronto, Toronto, Ontario, Canada; Hepatobiliary/Pancreatic Surgical Oncology Program, University Health Network, Toronto, Ontario, Canada

## Abstract

‘Early is too late’ describes a central problem in pancreatic cancer referring to its relentless drive to disseminate and highlights the need to understand how this disease spreads so rapidly. In this study, we profiled 1,013 samples from 277 donors, including tissue whole genome and RNA sequencing combined with plasma whole genome sequencing at ∼25x. Strikingly, local primary tumours, including some reaching 7cm, were found to shed little to no circulating tumour DNA (ctDNA). Instead, metastatic burden in the liver but not extrahepatic sites, was a main physiologic determinant of ctDNA levels. Whole-genome duplication (WGD), high cell cycle activity, and non-glandular differentiation emerged as tumour-intrinsic features related to increased ctDNA shedding. By contrast, decreased shedding was related to extrinsic features including a reactive microenvironment and unexpectedly, humoral immunity. Analysis of tumour clonal architecture showed that in patients with low ctDNA levels, the signal disproportionately originated from subclones and this signal persisted even when the primary tumour was removed. Disseminated subclones were a significant source of ctDNA in early-stage patients. Longitudinal analysis of patients revealed that subclones seeded metastases and shed ctDNA in the blood years before detection. This first report of paired tissue and plasma whole genomes in pancreatic cancer is a unique resource and has broad implications for disease surveillance, treatment monitoring, and early detection in this disease.

---

In many cancers, early-stage disease can be cured with surgery and adjuvant therapy, as tumour remains localized to the tissue of origin and micrometastases are chemosensitive. This is not true in pancreatic cancer (referring to adenocarcinoma) where removing local tumours often does not lead to cure. Survival rates for localized pancreatic cancer are dismal at 20 – 30%. Even in Stage I tumours, which are rare, distant metastases develop in half of patients. This has long supported the notion that primary and metastatic disease in most pancreatic cancer patients develops in parallel^1^. Regardless of the cancer type, detecting occult metastases is a challenge. Moreover, defining the window between local invasion and metastatic spread is key to developing effective screening strategies and understanding why early detection has worked in some cancers but not others.

To study this, we focused on circulating tumour DNA (ctDNA) as it provides a readout of cancer that is independent of tumour site. Traditionally, deep targeted sequencing has been used to detect low levels of ctDNA as ultra-high coverage is needed to capture rare mutations in the blood. However, this view changed when Zviran *et al.* (2020) showed that “mutation breadth” or tracking thousands of mutations with whole-genome sequencing (WGS), is more sensitive than deep targeted sequencing^2^. Although this approach is typically tumour-informed, plasma WGS can also be applied in a tumour-agnostic manner through the analysis of cell-free DNA (cfDNA) fragment features, a field known as fragmentomics^3–5^. Most studies using plasma WGS are generally performed in low-pass mode (<1x), and when higher depth plasma WGS (>20x) has been used, it has been on limited patient numbers and described in very few cancers^2,6,7^. To our knowledge, higher-depth paired plasma and tissue WGS studies across hundreds of patients has not been previously conducted in pancreatic cancer. Given the urgent needs of pancreatic cancer detection, we initiated a plasma WGS program to investigate ctDNA dynamics. Specifically, we set out to: (1) build a plasma WGS cohort from patients from all stages; (2) identify the determinants of ctDNA shedding in this disease; (3) assess whether plasma WGS can detect minimal residual disease (MRD) or subclinical metastases; and (4) evaluate its utility in treatment monitoring and progression.

## Genomic landscape of patient cohort

Accurate mutation tracking in the plasma requires matched mutation data from the tumour; however large patient cohorts where both plasma and tumour WGS have been banked are rare. We leveraged our longstanding biobank and trial infrastructure to assemble a patient cohort with matched tissue and plasma from all disease stages. Early-stage cases (Stage I/II) were obtained through surgical resections as part of International Cancer Genome Consortium (ICGC), and advanced cases (Stage III/IV) were collected from percutaneous biopsies through the COMPASS trial (Changes and Characteristics of Genes in Patients with Pancreatic Cancer for Better Treatment Selection; NCT02750657^8^). Among early-stage patients (n = 62), 47 plasma samples were collected prior to surgery (7 Stage I and 40 Stage II), with an additional 15 samples obtained after surgery (all Stage II) (**Supplementary Table 1-3**). In the advanced cohort, 12 patients had locally advanced disease and 130 had metastatic disease. All plasma samples from advanced stage patients were treatment naïve. Longitudinal sampling was carried out in 21 donors with 33 additional plasma draws at various points during their treatment course. For controls, plasma and matched germline DNA from peripheral blood were sequenced from 45 non-cancer individuals. In total, 305 plasma samples underwent WGS (depth: median 25x, range 10-72x). Together, plasma, tumour, and germline DNA WGS, along with tumour RNA-seq, comprised a resource of 1,013 samples from 277 donors.

Given the extremely low cellularity of pancreatic cancer, laser-capture microdissection (LCM) was performed on all tumour tissue (fresh-frozen) prior to WGS (n = 232 donors; median depth at 51x) (**Supplementary Table 4**). A median of 6,104 somatic single-nucleotide variants (SNVs), 451 small insertions/deletions (indels), and 103 structural variations (SVs) were observed per genome (**Supplementary Fig. 1a-c**). Mutation burden was higher in metastases than in primary tumours (p = 0.0048, < 0.001, 0.017 for SNVs, indels, and SVs, respectively). Recurrent somatic mutations were identified in classic driver genes, including *KRAS* (89%), *TP53* (83%), *CDKN2A* (81%), and *SMAD4* (44%) (**Supplementary Fig. 1d-g**). Eight percent of patients were homologous recombination deficient (HRD) (n = 18; **Supplementary Fig. 1h**), and two patients had mismatch repair deficiency (MMR) (**Supplementary Fig. 1i**). Copy-number profiling revealed recurrent arm-level gains in 1q, 8q and losses in 17p (**Supplementary Fig. 1j**), consistent with what is known in these known in these tumours. Whole-genome duplication (WGD) was present in ∼50% of tumours and was more recurrently observed in metastases compared to primary tumours (58% versus 42%; p = 0.0302) (**Supplementary Fig. 1k**). From paired RNA-seq data (n = 196), we also characterized our cohort by molecular subtype. Among primary tumours, 7% were Basal and 93% were Classical. In metastases, this shifted significantly to 25% Basal and 75% Classical (p = 0.001) (**Supplementary Fig. 1l**).

## When ctDNA are low, rare mutations paradoxically exist at high VAFs

We estimated cancer burden in plasma using a tumour-informed approach. To do this, we first applied MRDetect^2^, a highly sensitive method that uses a personal mutation profile from the tumour WGS and scans the paired plasma WGS for the same mutations. This approach allows simultaneous tracking of thousands of mutations and showed greater sensitivity for ctDNA detection than conventional mutation callers (**Supplementary Fig. 3a**). Using this, we were able to detect 1.4% to 91.6% (median = 10.3%) of tumour SNVs in the plasma across our cohort (**Fig. 1a**). Based on this, we stratified patients into three detection groups: low (0–10%), intermediate (10–50%), and high (50–100%). We first examined the cumulative variant allele frequency (cVAF) of each subgroup (**Fig. 1b**). cVAF is an accuate measure of ctDNA burden at low levels as it integrates the signal across all detected mutations. As anticipated, patients in the high-detection group showed high cVAFs and the intermediate-detection group showed lower VAFs. Strikingly, however, the cVAFs of patients in the low-detection group resembled that of the high subgroup. Moreover, the cVAFs of the low-detection group was significantly higher than in the intermediate subgroup (p = 0.00079). This was not a technical artifact as this pattern was absent in healthy controls (p = 0.11; **Supplementary Fig. 3b-c**). To understand this, we examined the distribution of individual VAFs among the groups and observed that the spike in cVAFs in the low-detection group was driven by individual SNVs that occurred at disproportionately high frequencies (Fig. 1c). This observation suggests that when ctDNA levels are low, the plasma signal is dominated by rare variants at high frequency and does not reflect the full mutation spectrum of the tumour.

**Fig. 1.**
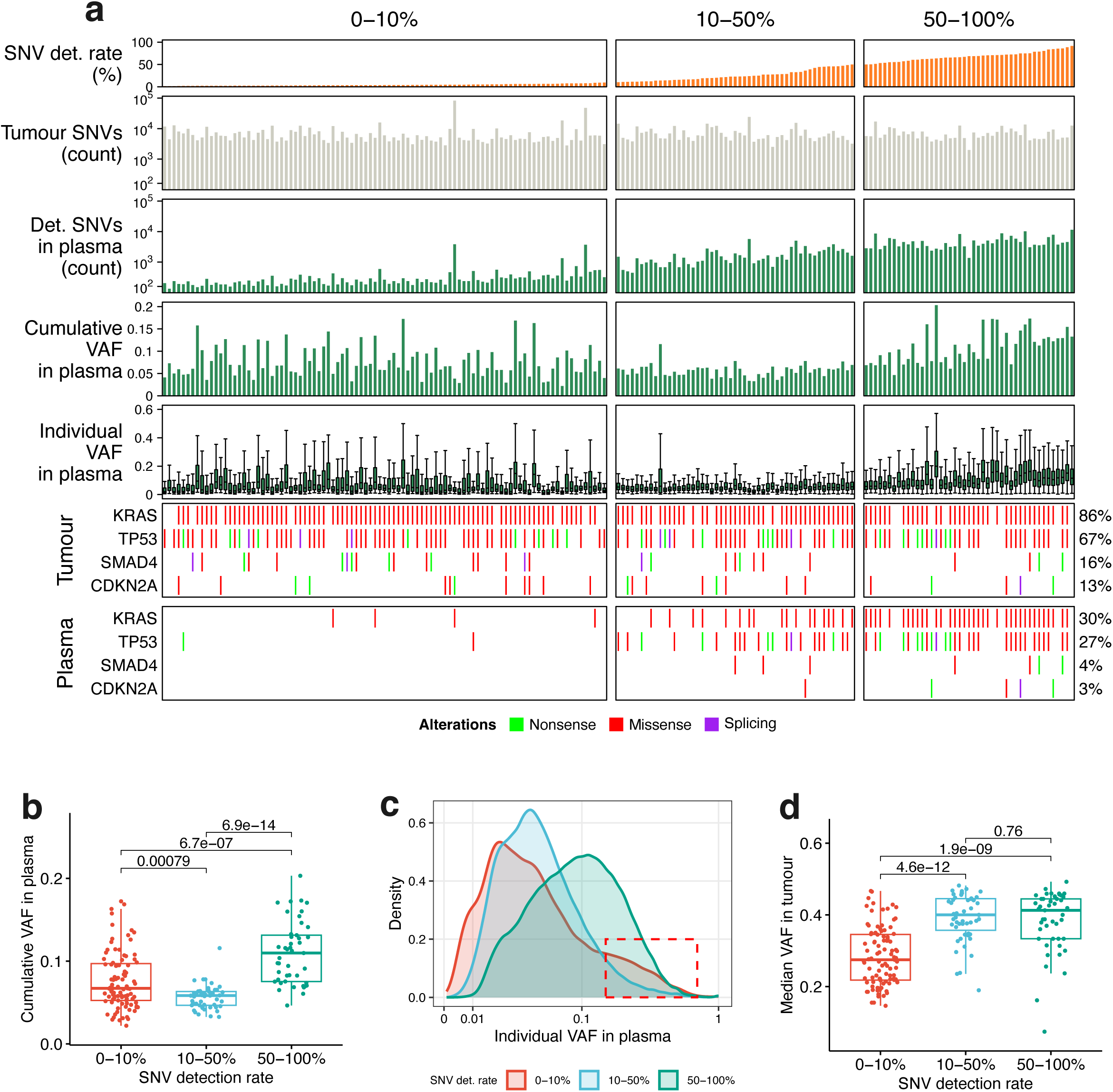
| Tumour-guided detection of mutations in treatment-naïve plasma. **a**, (1^st^ row) Detection rate was calculated by the number of SNVs detected in plasma over the number of total SNVs in the paired tumour WGS. Patients were stratified by detection rate. (2^nd^ row) The number of total SNVs in the paired tumour WGS. (3^rd^ row) The number of SNVs detected in plasma. (4^th^ row) Cumulative VAF of detected SNVs in plasma. (5^th^ row) The individual plasma VAFs of detected SNVs. The oncoprints for SNVs in 4 main drivers identified in paired tumour (6^th^ row) and plasma (7^th^ row). **b**, Cumulative plasma VAF of detected SNVs. **c**, Distribution of individual plasma VAFs of detected SNVs, stratified by detection rate. **d**, Median tumour VAFs of detected SNVs. The Wilcoxon signed-rank test was conducted between groups for **b** and **d**.

To understand this ‘low-burden, high signal’ paradox, we first examined whether these high-VAF mutations in the low subgroup corresponded to driver mutations (bottom, **Fig. 1a**). In the high-detection subgroup, driver mutations in genes such as *KRAS, CDKN2A, TP53*, or *SMAD4* were observed in the plasma in 84% of patients, and in 67% in the intermediate group. However, driver mutations were detected only in 6% of patients in the low-detection group. Thus, these high frequency SNVs in the low subgroup do not commonly reflect shedding of driver mutations. We then tracked these detected mutations for their VAFs in the paired tissue WGS. In the intermediate and high-detection groups, plasma SNVs corresponded to high VAFs in the tumour suggesting they were clonal in origin (**Fig. 1d**). However, in the low-detection group, these high VAF mutations had low VAFs in the tumour supporting they originated from subclones (p < 0.001, **Fig. 1d**). This supports that tumour clonal architecture impacts ctDNA dynamics and was investigated in great depth later in the study. These findings underscore the importance of genome-wide approaches at higher depth, which enable detection of signals that would have otherwise been missed by targeted or low-pass sequencing.

### ctDNA levels are uncoupled from primary tumour size and dependent on metastatic organ site

We developed a quantitative estimate of ctDNA burden that incorporates copy number and tumour cellularity. This method also corrects for the disproportionately high VAFs in the low-detection subgroup (**Supplementary Fig. 2, Methods**). We refer to this metric as plasma tumour fraction or TFx. TFx was validated through mixture analysis and benchmarked against established methods showing strong performance (**Supplementary Fig. 2c-d**). We also corrected TFx for background noise using healthy control samples (**Methods**). In simulations, TFx reliably detected ctDNA down to 5 × 10⁻⁴, given a typical tumour mutation load of ∼5,000 SNVs and a plasma sequencing depth of 20×–60× (**Supplementary Fig. 2c**).

We investigated ctDNA dynamics from different disease stages (**Fig. 2a**). After background correction, 97% of patients had detectable ctDNA (TFx > 0%; **Supplementary Fig. 4a**). However, the level of ctDNA was significantly different across stages. Among early-stage patients (Stage I/II, resectable tumours), the median plasma TFx was extremely low at 0.58% (range: 0% -5.88%) (**Fig. 2a**). We considered that Stage I patients would have lower plasma TFx compared to Stage II as these tumours are smaller and node-negative. However, this was not the case although our sample size is small (n = 7 in Stage I, p = 0.19) (**Supplementary Fig. 4b**). Among Stage II patients, node-positive patients trended toward higher plasma TFx compared to node-negative patients (0.73% versus 0.37%; p = 0.11, **Supplementary Fig. 4c**). Primary tumour size was similar between node-positive and node-negative patients (p = 0.54; **Supplementary Fig. 4d**). Overall, ctDNA was scarce in early-stage patients regardless of whether its Stage I or Stage II disease.

**Fig. 2.**
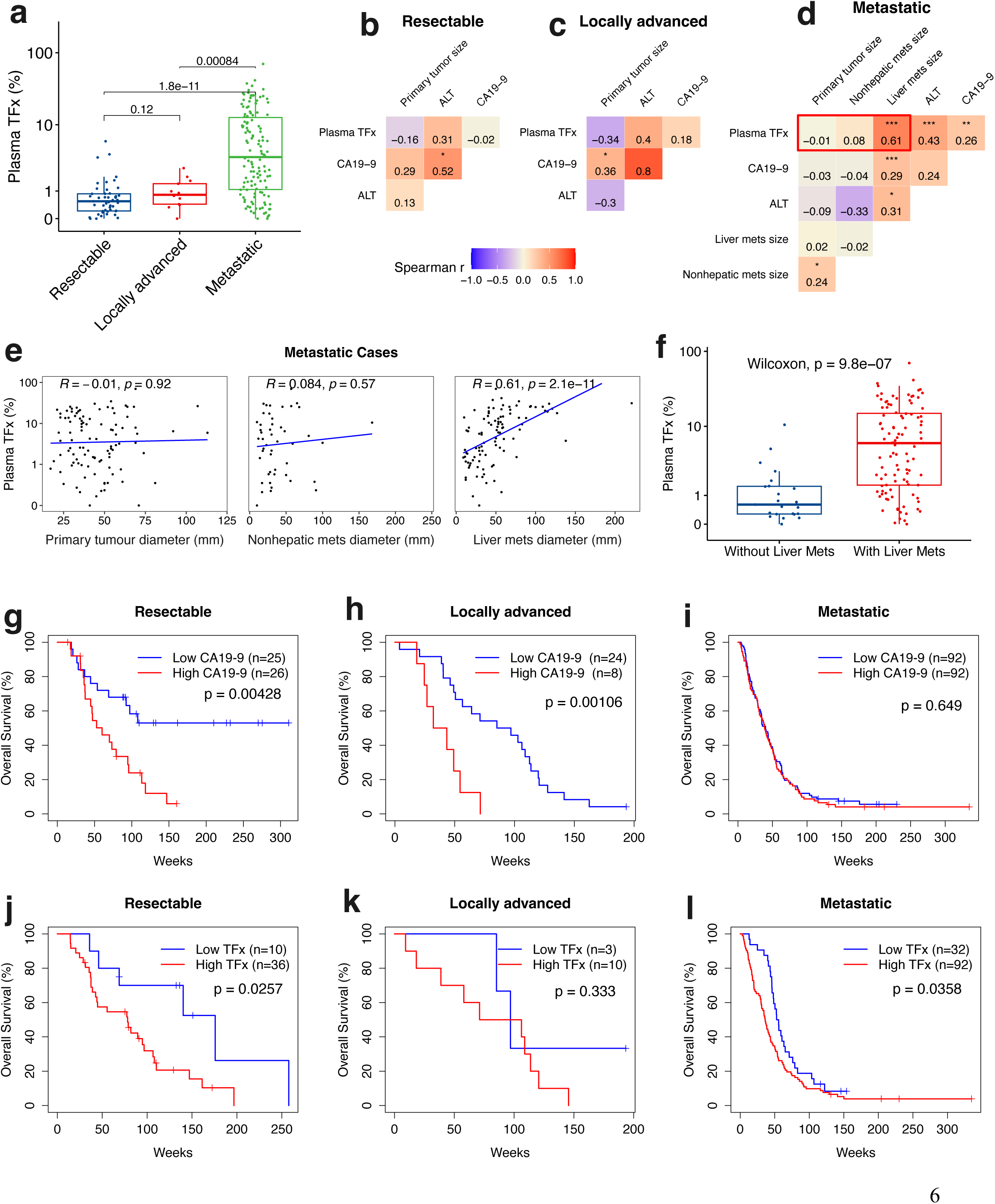
| The clinical associations of plasma TFx in pancreatic cancer. **a**, The levels of TFx in treatment-naïve plasma across different clinical stages of pancreatic cancer. The Wilcoxon signed-rank test was conducted. **b-d**, The Spearman correlations (shown in numeric values) between CA19-9, plasma TFx, ALT and tumour sizes across different clinical stages. *, p<0.05; **, p<0.01; ***, p<0.001. **e**, The scatterplots for plasma TFx versus sizes of primary tumour, nonhepatic and liver metastases from (**d**) labeled with a red box. **f**, Plasma TFx in Stage IV patients with and without liver metastases. The prognostic values of treatment-naïve CA19-9 (**g-i**) and plasma TFx (**j-l**) across clinical stages. The binary cut-off (low or high biomarker level) for optimal prognosis was determined per clinical stage using the Univariate Cox analysis (Supplementary Table 6). The optimal cut-off was then used to stratify the patients and the Kaplan-Meier analysis was used in the survival curves.

Patients with locally advanced pancreatic cancer (LAPC, Stage III) typically harbour larger primary tumours that are unresectable due to major blood vessel involvement. In LAPC patients, the median TFx was only slightly higher compared to early-stage patients (0.84% versus 0.58%; 1.4x; p = 0.12) (**Fig. 2a**). However, similar to early-stage patients, there was no correlation between ctDNA levels and size of the primary (**Fig. 2b, c**). To highlight this point, one LAPC patient had a large 7 cm primary tumour exhibiting a TFx of 0.45% (*Pcsi_1036*) (**Supplementary Fig. 7k**). These data show that even in LAPC where primary tumours were typically larger, ctDNA levels remained remarkably low. When disease was localized to the pancreas (Stage I-III), 77% of patients showed TFx below 1%.

In Stage IV metastatic disease, there was a dramatic rise in baseline plasma TFx to a median of 3.50% (range: 0%-69.80%) (**Fig. 2a**). This represents a 6-fold increase compared to early-stage patients (Stage I/II) and a 4-fold increase compared to LAPC patients. We first examined whether this increase in ctDNA levels was related to the size of the primary tumour in these patients. There was no relationship between size of the primary and ctDNA levels (R = -0.01, p = 0.92; **Fig. 2d, e**) and consistent with findings in localized disease. However, there was a strikingly strong correlation between TFx and size of the metastatic liver lesions. Although liver is the most common site of metastasis in pancreatic cancer, some patients present (18% of our cohort) with non-hepatic metastases (ex. lung or peritoneum). However, unlike liver metastasis, there was no correlation between size of non-hepatic metastases and TFx (R = 0.084, p = 0.57). Patients with non-hepatic metastases had markedly lower plasma TFx than those with liver metastases (p = 9.8 × 10^-7^; **Fig. 2f**).

Notably, 30% of metastatic patients showed TFx levels comparable to those in early-stage disease (<1% TFx) (**Fig. 2a**). This indicates that even after systemic dissemination, ctDNA levels remained low in a significant number of patients. Accordingly, low ctDNA levels need to be interpreted with caution as it may not reflect the presence or absence of metastatic disease. This has important implications for early detection and interpreting MRD dynamics. In summary, these findings support that ctDNA shedding in pancreatic cancer is dependent on organ-site.

## ctDNA and CA19-9 are not interchangeable clinical biomarkers

We benchmarked TFx against serum biomarker carbohydrate antigen 19-9 (CA19-9). CA19-9 data were available for a larger cohort (n = 299). As known, CA19-9 levels increased with clinical stage and mirrored the increase in ctDNA levels with progression (resectable: 210; LAPC: 344; and metastatic: 2,102 U/mL) (**Supplementary Fig. 5a**). However, when applying the standard CA19-9 positivity threshold (>37 U/mL), CA19-9 identified fewer patients than ctDNA (across stages: 78.4%-88.6% using CA19-9 versus 92.3%-98.5% using TFx) (**Supplementary Fig. 4a and 5b**). Five out of 134 (4%) patients were classified as CA19-9 ‘non-secretors’ (< 2 U/mL), and all had detectable ctDNA (TFx: 0.3%-15.2%) (**Supplementary Fig. 7a-c**). This is particularly useful in these patients as ctDNA is a tumour-specific signal in the absence of CA19-9.

Even though both CA19-9 and TFx increased with disease stage, they were poorly correlated (Stage I-III: no correlation; Stage IV: R = 0.26, p = 0.0084) (**Fig. 2b-d** and **Supplementary Fig. 7a-c**). There was also little to no correlation between CA19-9 and tumour size in the localized disease (**Fig. 2b, c**; **Supplementary Fig. 7d, e**). In metastatic patients, CA19-9 showed a weak correlation with liver lesion size (R = 0.29, p = 0.00032; **Fig. 2d** and **Supplementary Fig. 7g**), which is in stark contrast to the strong association found with ctDNA. The strength of CA19-9 correlation was similar to that of liver-function markers such as ALT, AST, and albumin (**Fig. 2d; Supplementary Fig. 6c**), supporting that CA19-9 reflects secondary effects of tumour burden in the liver. This is consistent with the known influence of CA19-9 on hepatic conditions such as cholestasis, biliary obstruction, stent irritation, and associated inflammation.

Next, we compared the prognostic performance of CA19-9 and plasma TFx. In resectable disease (Stage I/II), elevated pre-operative CA19-9 levels (p = 0.0043) and plasma TFx (p = 0.0257) were both prognostic (**Fig. 2g and 2j**). A decrease in CA19-9 post-operatively is a marker of good prognosis and this was consistent in our data (**Supplementary Fig. 5c, d**). However, we did not observe post-operative decline in ctDNA after removal of primary tumour in an unpaired analysis (p = 0.79) (**Supplementary Fig. 4e)**. In 14 out of 16 patients, TFx remained positive post-operatively. Even though ctDNA was detectable in most patients after surgery, patients with lower levels of ctDNA did show improved survival (**Supplementary Fig. 4f**). In metastatic patients, only ctDNA (p = 0.0358) but not CA19-9 (p = 0.649) was prognostic (**Fig. 2i and 2l**). These data have clinical implications as ctDNA is a tumour specific signal whereas CA19-9 reflects additional non-tumour related features. Importantly, these data indicate that ctDNA and CA19-9 are not interchangeable blood biomarkers.

## The influence of local and systemic effects on ctDNA shedding

Next, we examined molecular features related to ctDNA shedding. We used the metastatic cohort here as the dynamic range of ctDNA allowed us to separate low and high shedding patients, which was not possible in patients with local disease.

### **i.** Genome

From paired tissue WGS, we analyzed the impact of driver alterations on TFx. Mutations in *TP53*, *CDKN2A/B*, *SMAD4* or any specific *KRAS* mutant allele (G12D, G12V, G12R and others) showed no significant association with ctDNA levels (**Supplementary Fig. 8a, b**). There were several findings that were not significant but worth noting due to their translational importance. i) *KRAS* wild-type tumours trended to have higher TFx compared to *KRAS*-mutant tumours (**Supplementary Fig. 8a**); ii) In the *KRAS* mutated patients, copy-number imbalances favouring the mutant allele were related to higher TFx; iii) Patients with HRD showed modestly higher TFx than HR proficient patients (**Supplementary Fig. 8c**); and iv) Increased genomic instability as measured by structural variation^9^, tracked with higher TFx (**Supplementary Fig. 8c**). Among all variables analyzed, whole-genome duplication (WGD) was the sole genomic feature that was related to significantly higher TFx (p = 0.013) (**Fig. 3a**). Mechanistically, the elevated DNA content per tumour cell in WGD cases likely amplifies ctDNA signal upon cell death.

**Fig. 3.**
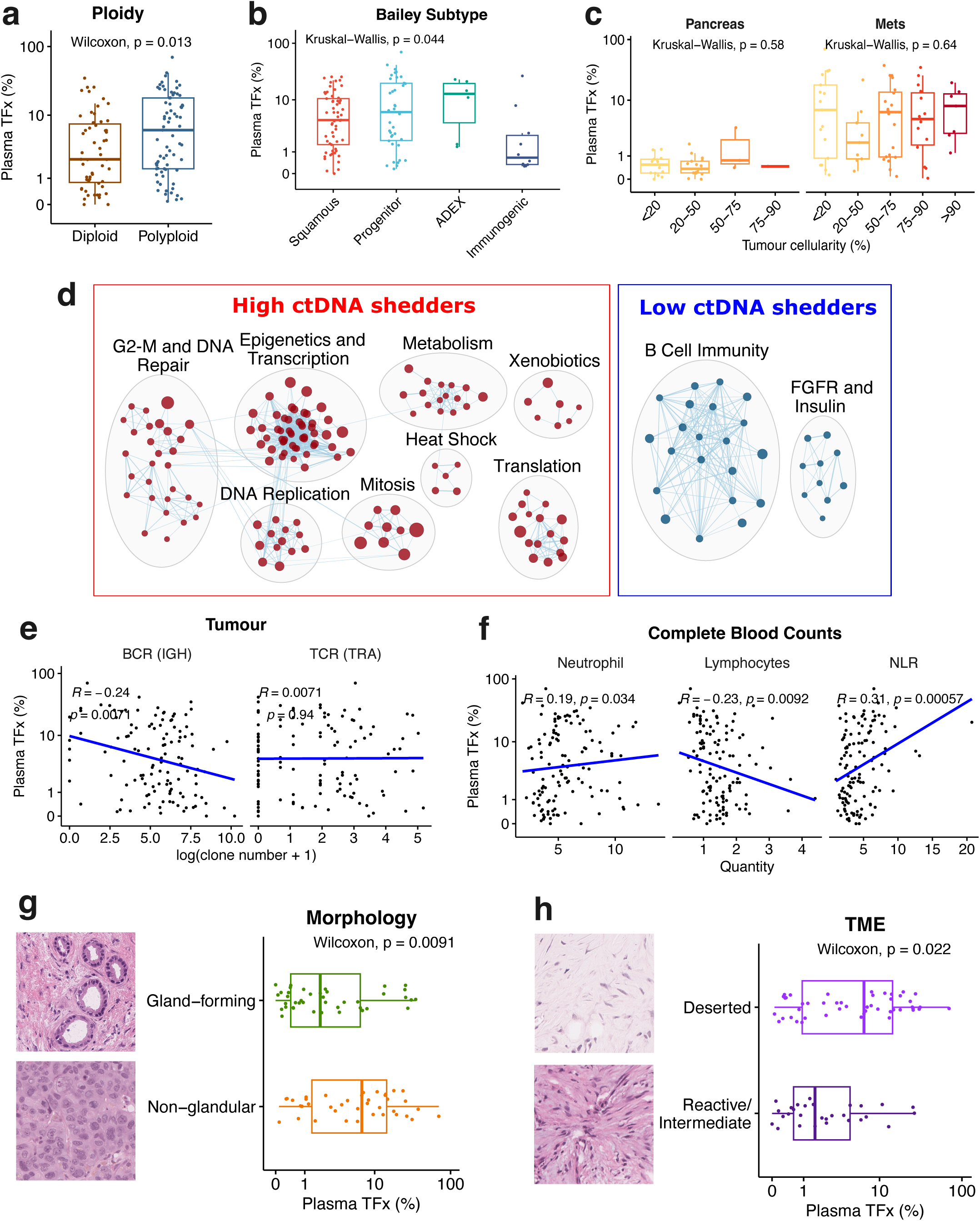
| Tumour intrinsic and extrinsic factors to influence ctDNA shedding in the metastatic pancreatic cancer. **a**, Plasma TFx in patients with diploid and polyploid tumours. **b**, Plasma TFx in patients stratified by tumour RNA subtypes from Bailey et al. 2016^10^. **c**, Plasma TFx versus histology-assessed tumour cellularity (pre-laser-micro-dissection) in primary pancreatic tumours and metastases. **d**, Enrichmentmap for the enriched Reactome gene sets in the paired tumours of high and low ctDNA shedders. The high and low shedders were stratified by using the median of plasma TFx in the Stage IV patients. **e**, The Spearman correlations between plasma TFx and immune repertoires in the tumour tissues. Repertoire abundance was defined as the number of detected clonotypes, using immunoglobulin heavy chain (IGH) for B cell receptors (BCR) and T cell receptor alpha (TRA) for T cell receptors (TCR), as identified by MiXCR. **f**, The Spearman correlations between plasma TFx and the quantities of neutrophils (×10^9^/L), lymphocytes (×10^9^/L) and the neutrophil-to-lymphocyte ratio (NLR) from the complete blood counts. **g**, Plasma TFx in patients with gland- and non-gland-forming tumour morphologies. Examples of the Hematoxylin and eosin (H&E) staining show a conventional (glandular) and a solid (non-glandular) tumour from liver biopsies. **h**, Plasma TFx in patients with deserted and intermediate/reactive tumour micro-environment (TME) (according to Grünwald et al. 2021^17^). All the analyses were conducted only in the Stage IV patients, except for (**c**).

### **ii.** Transcriptome and immune context

We extended our analysis to the transcriptome using paired RNA-seq data. Among the molecular classification schemes defined by several groups (Collisson *et al.*, Moffitt *et al.*, Bailey *et al.*, Ge *et al.*)^10–13^, only the ‘Immunogenic’ subtype showed an association with low TFx (**Fig. 3b**). The Immunogenic subtype is linked to tumour immunity, which we explored further through pathway analysis. Gene set enrichment (GSEA) between high-versus low-shedding tumours showed that high shedding tumours were enriched for cell-cycle and replication stress pathways (**Fig. 3d**), consistent with observations in other malignancies^14,15^. WGD tumours also exhibited increased cell cycle (**Supplementary Fig. 9a**), suggesting that increased proliferation of WGD tumours may also contribute to increased ctDNA release of these tumours. Low ctDNA shedding tumours showed strong enrichment for B-cell immunity signatures including B cell receptor (BCR) activation and related downstream Fc receptor–mediated activation and complement components (**Fig. 3d and Supplementary Table 7**). This was specific for B-cell programs as cytotoxic T-cell programs showed no differential enrichment. To exclude confounding by tumour cellularity, we inspected initial tumour specimens and found no association between cellularity and ctDNA levels (**Fig. 3c**). Because the enrichment of B-cell immunity was unexpected, we applied MiXCR^16^ to quantify immune receptor diversity from tumour RNA-seq. We found that increasing BCR (R = -0.24, p = 0.0071) clonotypes, but not T-cell receptor (TCR, R = 0.0071, p = 0.94), was related to decreased plasma TFx (**Fig. 3e** and **Supplementary Fig. 9b**). These data point to a potential role for humoral immunity in limiting ctDNA detection.

To extend this analysis to systemic immunity, we also examined complete blood counts. TFx correlated positively with circulating neutrophils (R = 0.19, p = 0.034) and negatively with lymphocytes (R = –0.23, p = 0.0092), resulting in an association with neutrophil-to-lymphocyte ratio (NLR; R = 0.31, p < 0.001) (**Fig. 3f**). Together, these findings suggest that infiltrating humoral immunity and systemic immune context (particularly B-cell function and neutrophil/lymphocyte balance) influence ctDNA dynamics.

### **iii.** Morphologic features and microenvironment

Morphologic analysis revealed that tumours with non-glandular features (cribriform, solid or squamous) shed significantly higher levels of ctDNA compared to glandular tumours (p = 0.009; **Fig. 3g**). We also examined the influence of the tumour microenvironment (TME) on ctDNA levels. We applied the Grünwald *et al.* framework^17^, which stratifies tumours into three microenvironmental states: (1) <u>Deserted</u> - immune-cold niches with lymphocyte depletion and myofibroblastic cancer-associated fibroblast (CAF) enrichment; (2) <u>Reactive</u> - immune-hot niches with abundant lymphoid and myeloid infiltrates; and (3) <u>Intermediate</u> - regions with mixed fibroblast populations and partial extracellular matrix (ECM) remodelling. Tumours with Deserted microenvironment exhibited significantly higher TFx than those with Reactive or Intermediate TMEs (p = 0.022) (**Fig. 3h**). Importantly, as stated above, these results are not confounded by tumour cellularity (**Fig. 3c**).

Overall, these results show that ctDNA shedding in pancreatic cancer is an interplay of tumour-intrinsic and microenvironment as well as local/systemic immunity.

## Fragmentomics uncovers a mutation-independent plasma signal in low-shedding disease

Mutation-informed assays will naturally miss ctDNA fragments that do not carry mutations. This is an important limitation to address especially when ctDNA is scarce. We analyzed the fragmentomic profile of our plasma cohort. ctDNA fragments tend to be shorter than cell-free DNA fragments from normal cells^18^. Thus, the ratio of short (100-150 bp) to long fragments (151-220 bp) (SLR) is an indicator of ctDNA quantity^3^. The sample-wise SLR was significantly higher in patients compared to controls and increased proportionally with TFx (**Fig. 4b and Supplementary Fig. 10a**). Crucially, even in the patients with extremely low plasma TFx (< 1%), SLR was significantly elevated compared to controls (p = 1 × 10^-10^).

**Fig. 4.**
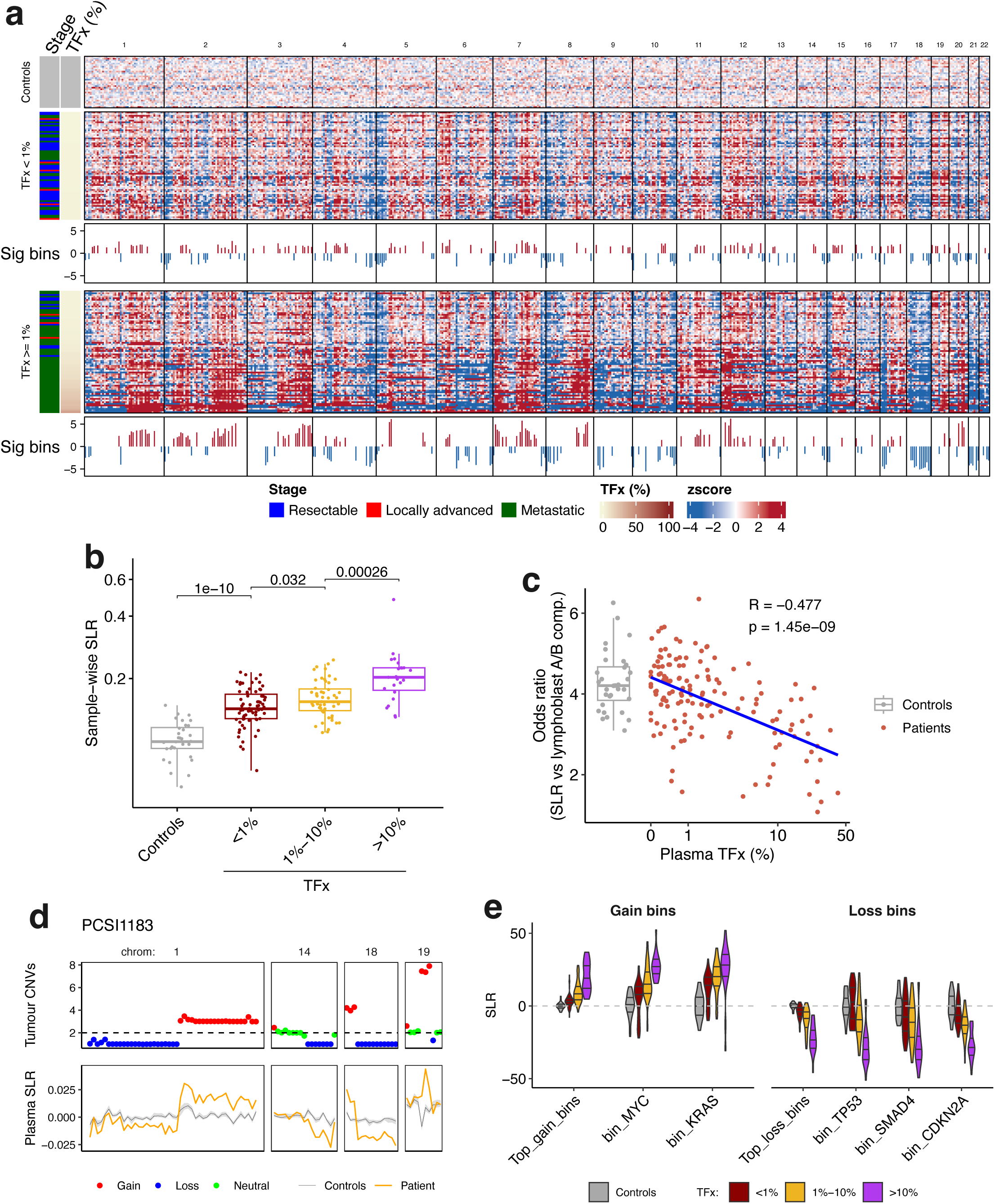
| cfDNA fragmentomics in plasma. Only treatment-naïve plasma samples were used from the patient cohort. **a**, The bin-wise short-to-long ratios (SLRs) in 512 non-overlapping 5 Mb genomic bins. SLR in each bin of every sample was z-normalized against 32 controls. The Wilcoxon signed-rank test was conducted in the unnormalized SLR between controls and patients, followed by the Bonferroni correction for multiple-testing. The significant bins with an adjusted p-value < 0.05 were labelled with the mean z score from the patients. **b**, Sample-wise SLR in the plasma from non-tumour controls and patients stratified by plasma TFx. The Wilcoxon signed-rank test was conducted between groups. **c**, The odds ratio between bin-wise SLRs and A/B compartments inferred from lymphoblast Hi-C data^19^. The Spearman correlation was conducted for the patients. **d**, An example (PCSI1183) to show the concordance between SLRs in plasma and CNVs in the paired tumour tissue in the corresponding 5 Mb genomic bins. A total copy number large than 2.3 and less than 1.7 was defined as copy number gain and loss, respectively. The gray line indicates the median and quantiles (5% and 95%) from 32 controls. **e**, Bin-wise SLRs in selected 5 Mb genomic bins in controls and patients stratified by plasma TFx levels. Top 50 bins with the most concurrent copy number gain or loss in tumours were selected.

We also examined the genome-wide fragment distribution patterns by calculating the SLR across the genome following the approach of Cristiano *et al.* (512 non-overlapping 5 megabase bins)^3^. Compared to controls, patients showed widespread deviations in bin-wise SLR across the genome, indicating that certain genomic segments were more affected by tumour-derived fragments (**Fig. 4a**). Because chromatin accessibility shapes nuclease cleavage, we compared bin-wise SLR to A/B compartments (open/closed chromatin) derived from Hi-C of lymphoblasts^19^, which is the main source of normal cfDNA in the plasma. As TFx increased, this correlation progressively weakened (R = –0.477, p < 0.001) (**Fig. 4c**), supporting a shift away from the hematopoietic chromatin landscape. In line with this, tumour copy-number alterations mirrored the fragmentomic landscape. Copy number gains in tumours corresponded to bins with higher SLR, whereas copy number losses showed the opposite effect (an example displayed in **Fig. 4d**). Canonical pancreatic cancer drivers followed this pattern (**Fig. 4e**). Amplified oncogenes such as *KRAS* and *MYC* were associated with higher SLR, while deleted tumour suppressors including *CDKN2A*, *SMAD4*, and *TP53* showed lower SLR. These loci exhibited dose-dependent fragmentomic shifts that scaled with TFx levels. A simple SLR-based classifier was able to distinguish patients from controls with an AUC of 0.96 at sequencing depth of 20x-60x for plasma WGS and still maintained great performance (AUC = 0.95) even at 0.5x (**Supplementary Fig. 10d, e**). Together, these findings indicate that the fragmentome produces a reproducible and mutation-independent signature of pancreatic cancer.

## Tumour subclones dominate ctDNA signal in low shedding disease

Next, we returned to earlier observations such as why large primary tumours shed very little ctDNA, and why patients with low ctDNA harboured mutations with high VAFs. The variables analyzed above could not explain these phenomena, which led us to examine the tumour clonal architecture. These analyses were possible as our paired tissue and plasma WGS were at a depth that enabled tracking of SNVs. This is one of largest analysis of its kind given the depth of the plasma and tissue genomes was performed across hundreds of patients. From tumour WGS, we classified SNVs as ‘clonal’ (present in all tumour cells) or ‘subclonal’ (restricted to a subset of tumour cells) based on cancer cell fraction estimates adjusted for copy number (**Supplementary Fig. 11; Methods**). In total, 1,445,734 clonal and 264,856 subclonal SNVs were identified (n = 238 patients). By tracking these mutations in the plasma, we observed a striking pattern. In high-shedding patients, clonal mutations overwhelmingly dominated the ctDNA signal (median clonal: 98%; subclonal: 2%; **Fig. 5a**). By contrast, in low-shedding patients, the contribution from clonal mutations declined sharply and there was a significant rise in the contribution from subclonal mutations (median clonal: 79%; subclonal: 21%; p = 5.4 x 10^-16^). Importantly, this effect was gradual, with the proportion of subclonal mutations progressively increasing as overall ctDNA levels declined. This was unexpected, since clonal mutations, which are present in all cells and significantly outnumber subclonal mutations, would be expected to dominate the plasma signal particularly when ctDNA levels are low.

As ctDNA levels are lowest in early-stage patients (**Fig. 2a**), we asked whether enrichment for subclonal ctDNA was more pronounced early in disease progression. Indeed, primary tumours showed a significantly greater contribution of subclonal mutations in the plasma compared to metastatic tumours (**Fig. 5b**), even when both groups were matched for ctDNA levels (**Fig. 5c**). The effect was most pronounced in Stage II primary tumours (**Fig. 5d**), which represent the main early-stage patient population. From tissue WGS, we confirmed that primary tumours harboured more subclonal mutations than metastatic tumours (**Fig. 5e**). This was expected as primary tumours have not undergone a bottleneck. However, even after accounting for this, the disproportionate representation of subclonal mutations under low ctDNA conditions persisted (**Fig. 5f**). Technically, this highlights that in tumour-informed assays, sampling the primary tumour to capture the full clonal diversity may be critical to capturing ctDNA shedding of rare subclones and early metastatic spread.

We sought to rigorously validate the above findings through a series of orthogonal tests. First, as noted above, accounting for increased number of subclonal mutations in primary tumours did not confound the analysis (**Supplementary Fig. 12a, b).** Second, we considered if stochastic sampling a small number of mutations would randomly enrich for rare subclonal mutations in ctDNA. Downsampling simulations showed that subclonal mutations are more difficult to capture when rare mutations are sampled indicating that our observation was not due to sampling bias (**Supplementary Fig. 13a**). We further performed random sampling tests to assess whether the selection of clonal versus subclonal mutations introduced any systematic bias. These analyses similarly ruled out the possibility of ascertainment bias (**Supplementary Fig. 13b**). Thus, this enrichment of subclonal mutations in low ctDNA conditions did not occur by chance. Third, we tested whether MRDetect introduced a bias in detecting subclonal mutations. This was possible as MRDetect scans billions of reads to identify rare mutations. To do this, we examined the distribution of mutant reads supporting clonal and subclonal mutations in both high- and low-shedding patients. In high-shedding patients, clonal mutations were supported by more reads, consistent with their higher abundance. By contrast, in low-shedding patients, mutant read distributions were equivalent between clonal and subclonal mutations, indicating that MRDetect did not preferentially detect subclonal variants under low ctDNA conditions (**p = 0.338; Supplementary Fig. 13c**). Finally, we developed an entirely independent approach to validate these findings using structural variants (SVs). SVs represent a different class of genomic alterations. By classifying SVs as clonal or subclonal (**Supplementary Fig. 14, 15** and **Methods**), we found that subclonal SVs were significantly enriched in patients with low shedding disease (p = 4.3x10^-9^; **Fig. 5g**). Together, these analyses establish that enrichment of subclonal mutations in low shedding pancreatic cancer patients is a genuine biological phenomenon and not a technical artifact. They also put into perspective our earlier findings. Low ctDNA levels in patient with large primary tumours occur as shedding in these patients likely reflects disseminated subclones. This work also explains why a subset of mutations in patients with low ctDNA levels exist at high VAFs in plasma as they likely represent disseminated cells (studied further below). Collectively, these findings demonstrate that ctDNA shedding in pancreatic cancer is shaped by the underlying clonal architecture of the tumour.

## Tracking patients with longitudinal sampling reveals pervasive early dissemination

Whether metastasis in pancreatic cancer is an ‘early’ or ‘late’ event is a controversial area and has profound implications for early detection ^20–23^. In ‘early’ dissemination, metastatic spread occurs in parallel with local invasion leaving little to no opportunity for curative intervention. In ‘late’ dissemination, there is a prolonged phase where the primary tumour is invasive but not metastatic, offering a window to cure the disease. The enrichment of subclonal mutations in the plasma in low shedding patients raised a natural hypothesis that this represents early dissemination. To investigate this, we assembled 21 patients with longitudinal sampling and detailed clinical annotations including 54 serial plasma timepoints and multi-tissue WGS in some cases. Follow-up ranged from 3 to 89 months. Our goal was to capture important clinical scenarios such as MRD and subclinical spread, early detection in high-risk individuals, and therapeutic resistance. All findings below were required to exceed background SNV detection rates from control samples to be significant (**Methods**).

### i. Subclones anticipate metastases

We first examined a patient who experienced rapid recurrence after resection (**Fig. 6a**). In patient Pcsi_0633, ctDNA was positive (TFx = 1%) three days after removal of the primary tumour with a persisting subclonal signal in the plasma. As the half-life of ctDNA is hours, this residual ctDNA could not have originated from the primary tumour. Fifty-seven days later, this patient developed liver recurrence, which was accompanied by a striking 37-fold spike in ctDNA (TFx = 37.4%). We performed WGS on both the primary and liver metastasis to track mutation dynamics. A subset of the subclonal mutations in the primary tumour had become fully clonal in the liver metastasis, which is strong evidence that persisting subclonal signal after surgery originated from disseminated cells that had previously seeded the liver (**Supplementary Fig. 16a**). The liver metastasis was whole-genome duplicated, whereas the primary tumour was diploid (**Supplementary Fig. 16b**). This is consistent with our findings that WGD is associated with increased ctDNA shedding. WGD is a hallmark of metastatic competence^24–28^. Remarkably, 54% (146/271) of plasma mutations three days post-surgery were unique to the liver metastasis when no metastases were radiographically visible (plasma sample 1 in **Supplementary Fig. 16c**). Moreover, private mutations from the liver metastasis significantly exceeded background detection rates (**Fig. 6a.ii**) and confirm active ctDNA shedding from occult micrometastases. This indicates micrometastases were actively shedding ctDNA long before clinical detection.

**Fig. 5.**
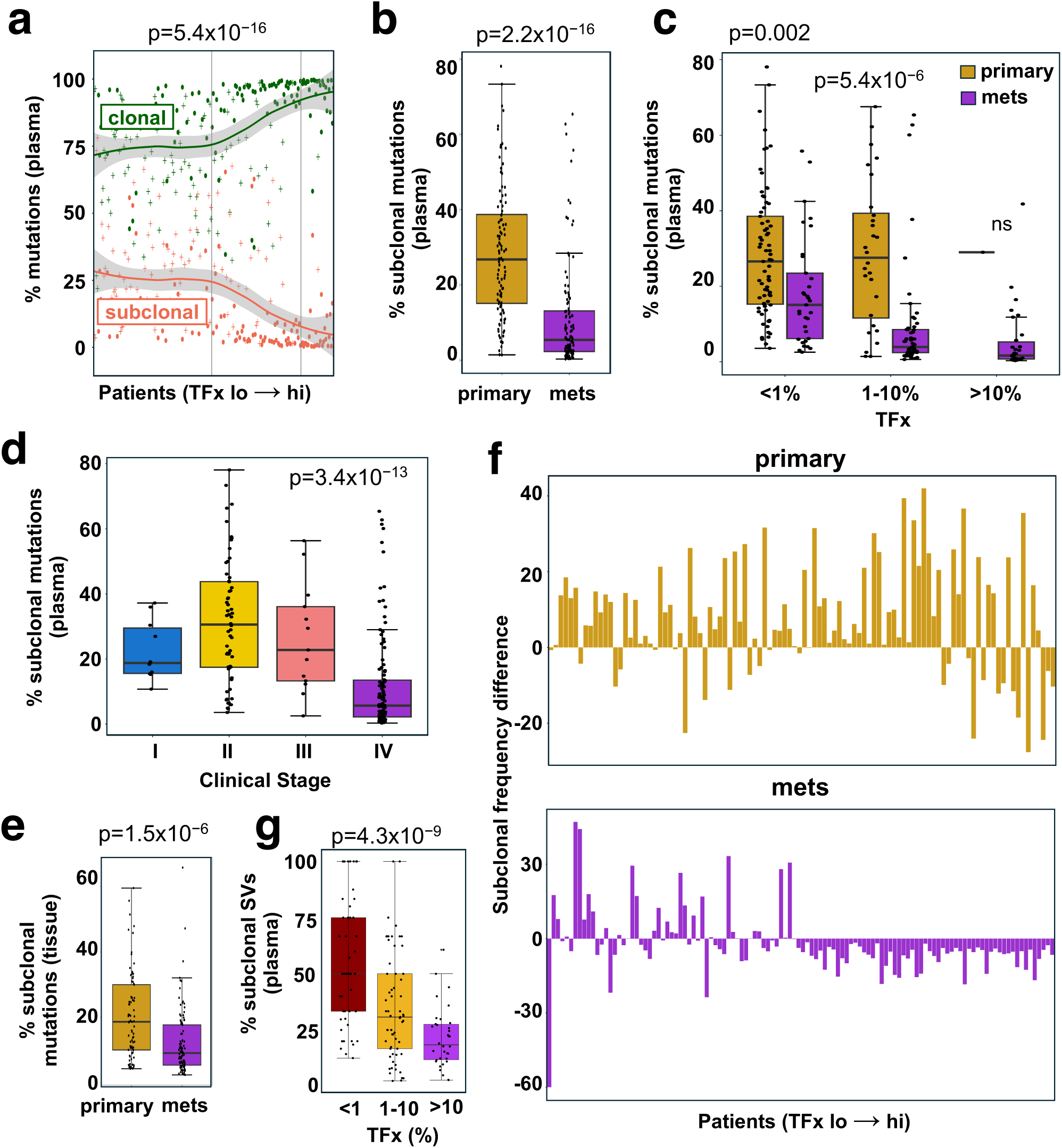
| Subclones contributed more to ctDNA at lower tumour fractions. **a**, Percent of clonal and subclonal plasma mutations across tumour fraction groups (plasma n = 218). Percent of subclonal plasma mutations between primary and metastasis samples (**b**), and stratified by TFx groups (**c**). A total of 218 plasma samples were involved. The Wilcoxon signed-rank test and the Kruskal-Wallis test were performed for **b** and **c** respectively. **d,** Percent subclonal plasma mutations across clinical stages. The Kruskal-Wallis test was performed (n = 198). **e,** Percent subclonal tissue mutations between primary and metastasis samples. The Wilcox test was performed (n = 174). **f,** The difference between the proportion of plasma subclonal SNVs and the proportion of bulk subclonal SNVs with increasing TFx (n = 218). **g,** Percent subclonal plasma SVs across TFx groups. The Kruskal-Wallis test was performed (n = 149).

**Fig. 6.**
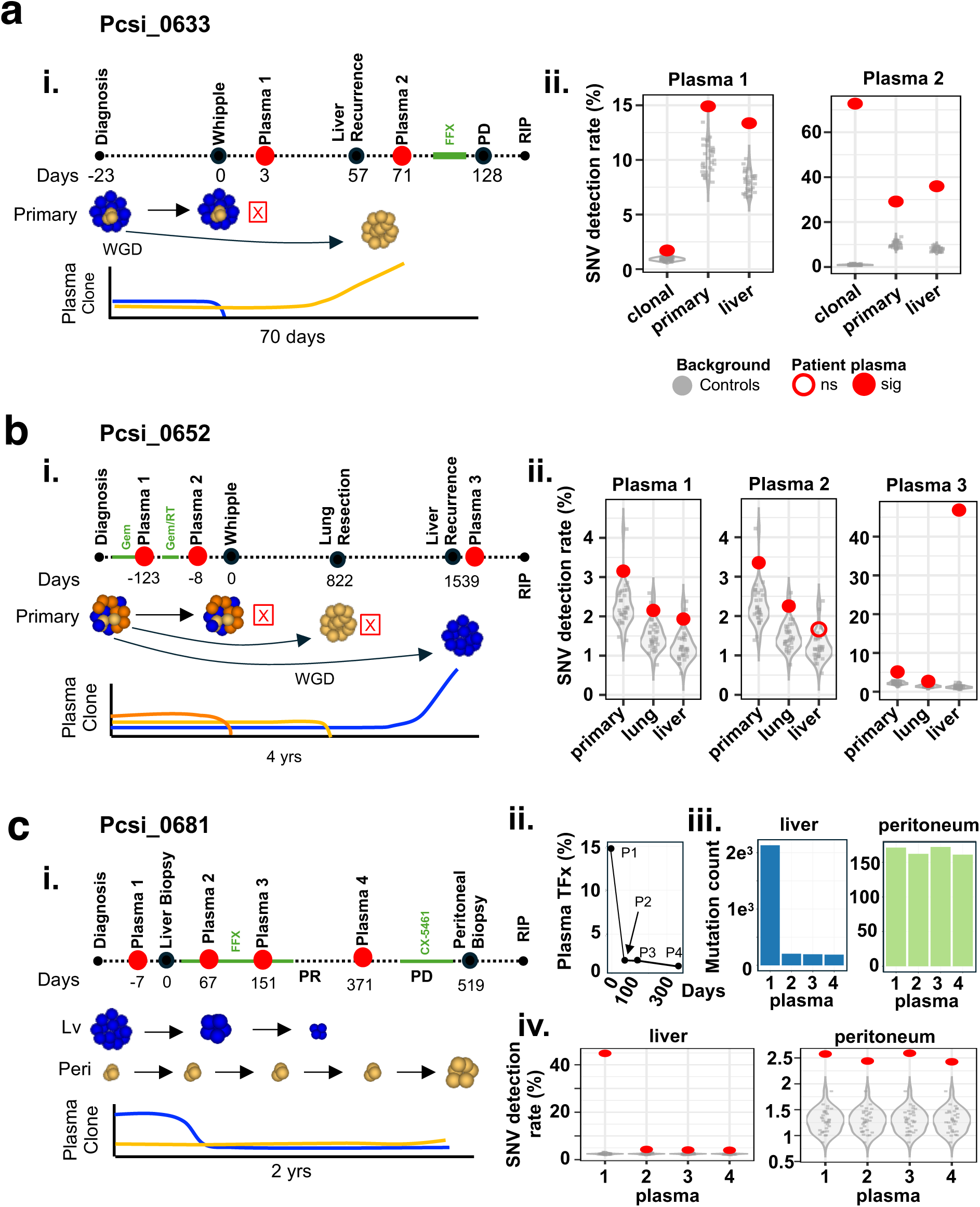
| Case studies with longitudinal plasma samples. **a**, PCSI0633 profile. i) Clinical timeline with schematic of cellular events. The green segment on the timeline indicates periods when the patient was on chemo- or radiotherapies. FFX, FOLFIRINOX; PD, progressive disease. ii) Detection rate of tissue-specific SNVs (# detected in plasma / total # in tissue). The background detection rate was calculated by searching tissue-specific SNVs in plasma of 32 healthy controls. The observed patient detection rate exceeding the 95% confidence interval of the background distribution (z > 1.645) was considered significant. **b**, PCSI0652 profile. i) Clinical timeline with schematic of cellular events. Gem, gemcitabine; RT, radiotherapy. ii) Detection rate of tissue-specific SNVs. **c**, PCSI0681 profile. i) Clinical timeline with schematic of cellular events. CX-5461, pidnarulex. ii) TFx at plasma timepoints. iii) The number of tissue-specific SNVs (liver and peritoneum) detected in plasma timepoints. iv) Detection rate of tissue-specific SNVs.

Similar MRD dynamics were observed in other patients. In Pcsi_0612, ctDNA remained detectable three days post-whipple and preceded radiographic evidence of an adrenal metastasis that appeared nearly a year later (**Supplementary Figs. 17a.i. and ii.**). In Pcsi_1056, plasma WGS detected ctDNA associated with a subclone that later gave rise to a brain metastasis more than a year later. (**Supplementary Figs. 17b.i. – iii**). Pcsi_1054 underwent a distal pancreatectomy to remove a tumour from the tail of the pancreas but remained positive for ctDNA post-operatively (TFx = 0.66%) (**Supplementary Fig. 17c.i. – iii**). Ten months later, this patient developed a recurrence in the head of the pancreas, followed six months later by a lymph-node metastasis. Mutations present in the recurrence in the head and the lymph-node metastasis were already detectable in the plasma 3 days post-operatively. Strikingly, ctDNA levels often remained relatively stable despite tumour growth and progression in many of these patients did show changes in site-specific shedding. Overall, tracking ctDNA dynamics of subclones can anticipate the emergence of metastases.

### ii. Detecting disease in high-risk individuals on molecular surveillance

As part of our longitudinal cases, there were two high-risk individuals with Peutz–Jeghers syndrome (PJS) (**Supplementary Fig. 18**). PJS is a rare, autosomal dominant cancer predisposition syndrome driven by germline loss-of-function mutations in *STK11* (*LKB1*). Both patients carried pathogenic *STK11* variants and ultimately developed pancreatic cancer arising from branch-duct intraductal papillary mucinous neoplasms (IPMNs)^29^. In Pcsi_0803, a 65-year-old woman, two plasma samples were collected, one at 6 months before diagnosis as part of surveillance, and another at diagnosis of the tumour (**Supplementary Fig. 18a.i**). WGS was also performed on tumour tissue obtained at diagnosis of her locally advanced cancer. Notably, ctDNA was already detectable 6 months before diagnosis, despite stable imaging of a 1.8 cm branch-duct IPMN. At diagnosis, there was a modest rise in ctDNA levels, and 20-25% mutations detected corresponded to a tumour subclone (**Supplementary Fig. 18a. ii** and **iii**). Pcsi_0666 was a 38-year-old male carrier who was diagnosed with metastatic pancreatic cancer (**Supplementary Fig. 18b.i**). Three plasma draws were conducted, one collected 18 months before diagnosis during surveillance, one at diagnosis, and another during treatment. ctDNA was already detectable 18 months before diagnosis where imaging showed only a 0.7-cm branch-duct IPMN. An important limitation to acknowledge is that we cannot exclude the possibility that the ctDNA originated from the IPMN rather than the invasive cancer. While this possibility cannot be excluded, the mutations tracked in this patient were derived from the liver metastasis (**Supplementary Fig. 18b.ii**), supporting that dissemination had already occurred at the time of surveillance in this second patient. Moreover, a rare SV unique to the metastasis was also detected in the surveillance plasma sample (**Supplementary Fig. 18b.iii**), indicating that chromosomal instability had already been established by this stage. These findings underscore the importance of molecular surveillance in addition to imaging in high-risk individuals.

### iii. Tracking of latent metastatic lineages

Approximately 16% of pancreatic cancer patients develop lung metastases without liver involvement^30^. This metastatic pattern is associated with a more indolent course and improved survival, but its evolutionary origins are unknown. Patient Pcsi_0652 initially presented with local disease that was surgically removed and two years later developed a lung metastasis (**Fig. 6b i**). The lung lesion was resected, and the patient remained disease-free for another two years before developing a liver metastasis soon after which the patient passed away. Over this four-year period, we obtained tissue from the primary tumour, the lung and liver metastasis, and performed WGS across all tissues to reconstruct tumour development. Although we initially presumed that lung metastasis gave rise to the liver metastasis as the primary tumour was removed 4 years prior, several lines of evidence indicated that the primary seeded both metastases independently. First, the primary tumour shared distinct mutation clusters with each metastasis but there were few to no mutations shared between the two metastases themselves (**Supplementary Fig. 19a**). Second, despite the four-year interval between removal of the primary tumour and the emergence of the liver metastasis, these two were more closely related, sharing 1,034 SNVs, whereas only 100 mutations were shared between the primary and the lung metastasis (**Supplementary Fig. 19b**). Third, a loss-of-heterozygosity (LOH) event on chromosome 22 was observed exclusively in the lung metastasis, while the primary and liver lesions retained both parental alleles (**Supplementary Fig. 19c**). Because a lost chromosome cannot be regained, this is strong evidence that the lung did not seed the liver. Together, this indicates that the both the lung and liver metastases were seeded independently by the primary tumour years before clinical detection. Strikingly, private mutations to the lung and liver metastasis were detectable in the plasma collected at diagnosis, and years before they would emerge (**Fig. 6b ii**). A second patient, Pcsi_0473, exhibited a similar pattern of early dissemination (**Supplementary Fig. 20a**). This patient experienced multiple lung recurrences following resection of the primary tumour, with four plasma samples spanning 7.5 years. ctDNA levels rose modestly before the third lung recurrence and remained persistently detectable throughout the disease course (0.4%, 0.3%, 0.8% and 0.7% across seven years). Although only the primary and final lung metastasis was sequenced, private mutations from the lung lesion were found in the earliest plasma sample, indicating that the metastatic lung lineage had disseminated at diagnosis and was actively shedding throughout this patient’s treatment journey (**Supplementary Fig. 20b**). Importantly, the majority of mutations in this patient derived from subclones supporting that tumour had disseminated before diagnosis (**Supplementary Fig. 20c**). This demonstrates that dissemination occurred years before metastases became visible.

### iii. Therapeutic evolution and organ site specific responses

Although many of the patients above received therapy during their clinical course, one case warrants particular attention due to its therapeutic relevance. Pcsi_0681 harboured a germline *BRCA2* frameshift mutation with a second hit due to LOH on chromosome 13, which conferred HRD. This patient presented with advanced disease involving both liver and peritoneal metastases (**Fig. 6c i**). Tissue from both metastatic sites and 4 serial plasma samples were collected over approximately a 2-year period. Consistent with HRD, this patient showed a dramatic response to platinum-based therapy. Plasma TFx dropped from 14.8% to 1.29% following treatment with FOLFIRINOX (**Fig. 6c ii**). From the tissue WGS, we were able to track site specific mutations from the liver and the peritoneal metastasis. The number of detected liver-specific mutations declined rapidly from 2,132 (plasma 1) to ∼200 (plasma 2-4), while peritoneal-specific variants stayed constant (∼160; plasma 1-4) (**Fig. 6c.iii)**. This suggests that the dramatic treatment response in this patient predominately derived from the liver metastasis. Notably, a *BRCA2* frameshift reversion mutation associated with platinum resistance emerged from the peritoneal lesion (**Supplementary Fig. 21**). This reversion mutation aligned to both the reduced sensitivity to therapy of this lesion and no change in the number of peritoneum-specific mutations detected in plasma over time.

Collectively, these longitudinal data provide a unique window into the clonal dynamics of pancreatic cancer progression, revealing that metastatic dissemination is established months to years before it becomes radiographically detectable.

## Discussion

Our findings provide a molecular explanation for why “early is too late” in pancreatic cancer. Large, bulky primaries shed little ctDNA in early-stage patients and often hover at ctDNA detection limits. Two factors likely underpin this low shedding in localized disease. First, primary pancreatic tumours are embedded in dense stroma, which likely impacts apoptotic release of ctDNA. Second, and perhaps more clinically relevant, is that in early disease, a significant part of the ctDNA signal originates from disseminated subclones rather than the primary tumour. This explains the persistence of ctDNA after surgical resection and also contributes to the overall low shedding given that these micrometastases are not detectable. Notably, we also found that ctDNA shedding is also organ-site dependent. Hepatic metastatic burden is the main driver of the ctDNA, whereas extrahepatic sites contribute less. It is noteworthy that nearly a third of metastatic patients still shed at levels comparable to localized disease, suggesting pancreatic cancer is not a high ctDNA shedding disease. We also found that both tumour-intrinsic (e.g., genome doubling, cell-cycle activity) and tumour-extrinsic factors (immune context, stromal reactivity) influence ctDNA dynamics, indicating that shedding reflects the biological state of the tumour rather than just size. Together, these findings suggest that ctDNA is not just a biomarker of the presence of the tumour but a dynamic readout of tumour physiology and evolution. These are important considerations in cancer detection, monitoring and interception.

Our integrating paired tissue and plasma was fundamental to linking genomic architecture with plasma dynamics. However, we recognize that this is tumour-informed and not always feasible particularly in the context of early detection or population screening. In the future, a combined strategy will likely be needed. Tumour-informed approaches such paired tissue and plasma WGS are uniquely positioned to track dissemination, evolution, and therapeutic response, whereas tumour-agnostic assays (e.g., fragmentomics and other emerging multi-feature classifiers) are better positioned for broader detection when tissue is unavailable^31–33^. Regardless of the approach, our findings emphasize that genome-wide resolution is essential, as much of the biological signal at very low ctDNA fractions will be missed by targeted panels. Pancreatic cancer’s defining feature is its capacity to disseminate early and develop rapidly. To intercept this disease, detection sensitivity must extend to the level of precursor lesions. The IPMN cases in our cohort suggest this is achievable, though identifying PanIN, the main precursor of this disease, poses a greater challenge. Importantly, recent work by Braxton *et al.* demonstrates that PanIN lesions can extend to centimetre scale^34^, raising the possibility that these early precursors may be detectable in plasma as they expand. Ultimately, meaningful progress will depend on shifting intervention earlier in the disease trajectory. Only by intercepting this cancer before it disseminates can we hope to significantly alter the course of this disease.

## Methods

### Patients with pancreatic cancer and non-tumour controls

The current study cohort consisted of patients with pancreatic cancer (pancreatic ductal adenocarcinoma, PDA) across all clinical stages (Stage I-IV) who had tumor tissue and/or plasma cfDNA sequencing (a detailed summary is provided in **Supplementary Table 1**). Written informed consent was obtained from all patients. Most of the patient were enrolled at Princess Margaret Cancer Centre at the University Health Network (UHN), Toronto, Canada. Most of the resectable and advanced cases were obtained through the International Cancer Genome Consortium (ICGC) and the COMPASS trial (no. NCT02750657), respectively, with some additional ones through the Hepato-Pancreato-Biliary (HPB) Tumour Tissue Banking. Tumour or plasma samples of patients were banked at either the UHN Biospecimens Program (Biobank) or the Ontario Pancreas Cancer Study (OPCS). Plasma samples of non-tumour controls were collected through and banked at OPCS. Approval for the study was obtained through the University Health Network Research Ethics Board (nos. 03-0049, 08-0767, 15-9596, and 20-5594). A subset of the cohort was obtained from Sunnybrook Health Sciences Centre (Toronto, Canada), Kingston General Hospital (Kingston, Canada), McGill University (Montreal, Canada), Mayo Clinic (Rochester, the USA), Massachusetts General Hospital (Boston, the USA) and Sheba Medical Centre (Tel Aviv, Israel)^22,35–37^.

### Tumour tissue processing

For resectable cases, tumour tissues were obtained from surgical specimens. For locally advanced and metastatic cases, tumour tissues were obtained by image-guided percutaneous core needle biopsy. Fresh tumour tissues were embedded in optimal-cutting-temperature (OCT) compound and snap-frozen in liquid nitrogen. For tumour enrichment, laser capture microdissection (LCM) was performed on the frozen tumour tissues.

### Tumour and normal DNA Whole-genome Sequencing

Whole-genome sequencing (WGS) was conducted as described in the previous studies^22,35^. DNA was extracted from LCM’d tumour tissues and normal tissues using Gentra Puregene Tissue Kit components, while DNA was extracted from buffy coat using the Gentra Puregene Blood Kit. The Qubit dsDNA High Sensitivity kit was used to quantify the extracted DNA. Illumina paired-end libraries were prepared using either the NEBNext DNA Sample Prep Master Mix Set, the Nextera DNA Sample Prep Kit or KAPA Library Preparation Kits. Libraries were quantified on the Illumina Eco Real-Time PCR Instrument using KAPA Illumina Library Quantification Kits. Then, paired-end sequencing was conducted on the Illumina HiSeq 2000/2500 platform using 2 × 101 cycles with TruSeq Cluster kit v.3 (Illumina, no. PE-401-3001/FC401-3001) and 2 × 126 cycles with HiSeq Cluster kit v.4 (Illumina, no. PE401-4001/FC-401-4001), combined with rapid-run (3) 2 × 101 cycles HiSeq SBS kits (Illumina, no. PE-401-4002/FC-402-4023), or the NovaSeq 6000 (2x151 or 2x101 cycles). Sequencing coverage was targeted at 50× and 35× for tumour tissues and normal DNA, respectively.

### Tumour genomic alternation profiling

Raw sequencing reads were aligned to the human reference genome (hg38) using Burrow-Wheeler Aligner (BWA, version 0.7.17)^38^. Germline variant calling was performed using Genome Analysis Tool Kit (GATK, version 4.1.2)^39^. Somatic single nucleotide variants (SNVs) were identified as the intersection of calls by Strelka2 (version 2.9.10)^40^ and MuTect2 (version 4.1.2)^41^. Small somatic insertions and deletions (INDELs) were identified using the consensus between two of four callers: Strelka2, MuTect2, SVaBA (version 134)^42^ and DELLY2 (version 0.8.1)^43^. Somatic structural variants (SVs) were identified using the consensus of 2 of 3 callers: SVaBA, DELLY2, and Manta (version 1.6.0)^44^. HMMcopy (version 0.1.1) (https://github.com/shahcompbio/hmmcopy_utils) and CELLULOID^22^ were used to call copy number segments, tumour cellularity, and ploidy.

### Tumour RNA Sequencing

RNA was isolated from LCM’d tumour tissues using the PicoPure RNA Isolation Kit. RNA was treated with the RNase-free DNase Set and quantified using the Qubit dsRNA High Sensitivity kit. RNA quality was measured using both the RNA Screen Tape Assay and 2200 TapeStation Nucleic Acid System. Only RNA with RNA integrity number (RIN) scores > 7 was used for library preparation. Complementary DNA libraries were prepared using the TruSeq RNA Access Library Sample prep kit. Library pools were quantified on the Eco Real-Time PCR Instrument using KAPA Illumina Library Quantification Kits. Paired-end sequencing of 2 × 126 cycles was conducted on the Illumina HiSeq 2500 platform with a target of minimally 30 million unique mapped reads. Reads were aligned to the human reference genome (hg38) and transcriptome (Ensembl v.100) using STAR v.2.7.4a^45^.

### Enrichment analysis

Raw counts of all the genes from tumour RNA-seq were obtained by HTSeq (version 2.0.4)^46^, and were used for differential expression analysis between two groups by DESeq2 (version 1.30.1)^47^. Genes were sorted by log2FoldChange and were used for Gene Set Enrichment Analysis (GSEA)^48^, which was conducted by clusterProfiler (version 4.14.4)^49^. Gene sets, including Hallmark^50^ and Reactome^51^ were used in the analysis. The GSEA results were visualized using Cytoscape^52^.

### MiXCR for immune profiling

MiXCR (v3.0.13)^16^ was used to profile the clonotypes of T and B cell receptor (TCR and BCR) from tumour tissue RNA-seq. The “*analyze*” module was used with the following parameters: “*shotgun --species hsa --starting-material rna --only-productive*”.

### Tumour genomic and transcriptomic features

Patients’ tumour samples were classified as homologous recombination deficiency (HRD) using the hallmarks of HRD, as described in the previous publications^53,54^. The HRD hallmarks include hits in HRD genes, SNV and SV counts, 4 bp+ deletion, 100-10kbp deletion, and 10k-1mbp duplication loads, C to T ratio, 4 bp+ deletion ratio and 100-10kbp deletion ratio. Tumours with at least 6 of these hallmarks were defined as HRD, and the ones with 3-5 hallmarks were manually curated.

Tumours were classified as DNA mismatch repair (MMR) deficient if any biallelic pathogenic variant was found in one of the MMR genes (*MLH1*, *MSH2*, *MSH6*, *PMS2* and *EPCAM*) from tumour WGS^55^.

The *KRAS* subtypes were defined according to the previous work^8^. The tumours with no nonsynonymous SNV in *KRAS* were wild type (WT). In the *KRAS* mutant tumours, the ratio of the mutant copy number to the WT copy number in the *KRAS* gene was calculated. The balanced, minor imbalanced and major imbalanced classes were the tumours with a ratio of 1, 1-2 and ≥3, respectively.

All the other genomic and transcriptomic subtypes in PDA were determined using either tumour WGS or RNA-seq according to the classifications in previous publications: Waddell^9^, Moffitt^13,56^, Ge^12^, Collisson^57^ and Bailey^10^.

### Plasma collection and cfDNA extraction

Blood samples were collected from patients diagnosed with pancreatic cancer and from healthy individuals under approved institutional ethics protocols. Blood was drawn into EDTA tubes, kept on ice and processed within 4 hours of blood collection. Plasma was separated by centrifugation at 1,500 x g for 20 minutes at 4 °C, followed by a second centrifugation at 16,000 x g for 10 minutes to remove residual cellular debris. Plasma aliquots were stored at -80 °C until cell-free DNA (cfDNA) extraction.

For plasma received from OPCS, samples were collected in ACD tubes and processed within 24 hours by centrifugation at 1,000 x g for 10 minutes. cfDNA was extracted from 1-5 mL of plasma using the QIAamp Circulating Nucleic Acid Kit (Qiagen) or the KingFisher MagMAX Cell-Free DNA Isolation Kit (Thermo Fisher Scientific) according to the manufacturer’s instructions. DNA concentrations were assessed using the Qubit dsDNA HS Assay Kit (Thermo Fisher Scientific).

### Plasma whole-genome sequencing

Plasma WGS libraries were prepared using the NEBNext Ultra II DNA Library Prep Kit for Illumina (New England Biolabs). Fragmentation was not performed for the majority of the samples due to the naturally short size of cfDNA. The sheared-fragment libraries were not included in the fragmentomic analyses (see below). End-repair, A-tailing, and adapter ligation were performed on an input of 5-40ng cfDNA following the kit protocol. Libraries were amplified using 4-6 cycles of PCR and purified using SPRIselect beads (Beckman Coulter). Library quality and size distribution were verified using the Bioanalyzer or Fragment Analyzer (Agilent Technologies). Libraries were sequenced on an Illumina HiSeqX, Illumina NovaSeq 6000 (for most samples) or Illumina NovaSeq X Plus platform using 150 bp paired-end reads, targeting a mean coverage of 30-60x across the genome. Sequencing data were demultiplexed and converted to FASTQ format using Illumina’s bcl2fastq software. Read alignment and standard conventional somatic mutation calling were performed using the same approaches as the tumour tissue WGS described above.

### Plasma tumour fraction estimation using ichorCNA

ichorCNA (version 0.2.0)^58^ is a tumour-agnostic approach to estimate plasma tumour fraction using copy number variations in the plasma. A bin size of 500 kb was used for the current plasma cohort. A panel of normal was generated from 10 randomly selected healthy plasma WGS from the current cohort and was provided to *--normalPanel*. Following parameters were used to account for lower tumour fraction samples: *--estimateScPrevalence FALSE --chrs "c(1:22) " --normal "c(0.5, 0.6, 0.7, 0.8, 0.9, 0.95, 0.99, 0.995, 0.999)"*, while all the other parameters were maintained as default.

### Tumor-guided SNV calling in plasma using MRDetect

A tumour-guided approach, MRDetect^2^, was used to identify SNVs in patients’ plasma WGS. Briefly, a list of somatic SNVs discovered from the tumour WGS (as described above) was provided to MRDetect, and then it searched the corresponding loci in the paired plasma WGS BAM/CRAM file to screen for the mutant reads. MRDetect applied a trained support vector machine (SVM) to filter out mutant reads that were potentially sequencing artifacts to increase the accuracy of the SNV calling. As patient plasma usually has a low tumour fraction, most of the true SNVs in the plasma can have very low variant allele frequencies (VAFs) and can be easily removed by the conventional locus-based variant callers due to low supporting mutant reads. MRDetect is read-based and it can largely rescue the low VAF SNVs that can be potentially filtered by the conventional variant callers and hence increase the sensitivity of SNV calling in plasma. The evidence of SNVs in the tumour indicates that the detection of even only a few mutant reads in the paired plasma is likely true positive.

### Plasma tumour fraction estimation using a Bernoulli model

Following SNV calling in plasma by MRDetect, Zviran et al. 2020 used a formula based on the Bernoulli model to estimate plasma tumour fraction^2^:

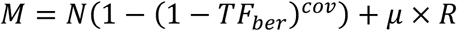

where M is the number of SNVs detected in plasma, N is the number of SNVs in the paried tumour, TF_ber_ is the Bernoulli TFx, cov is the local coverage at the loci with a tumor-specific SNV, µ is the noise rate (calculated by the number of SNVs detected in control plasma over total reads checked) and R is the total number of reads checked over the loci.

### Plasma tumour fraction estimation using a novel algorithm

We developed a tumor-guided algorithm to estimate plasma tumour fraction (denoted as TFx), following plasma SNV calling by MRDetect. For the i^th^ SNV locus in a patient’s tumour WGS, the VAF can be calculated as below:

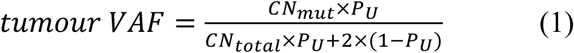

where CN_mut_ is the mutant copy number of this SNV in the locus, P_U_ is the tumour purity, CN_total_ is the total copy number in the locus in the tumour. The tumour SNV VAF, P_U_ and CN_total_ can all be calculated from tumor WGS. In the current study, the VAFs of SNVs were derived from Strelka2, while P_U_ and CN_total_ were calculated by CELLULOID. Then, the equation can be solved for CN_mut_:

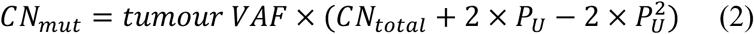

When the tumour cells break and release ctDNA into the blood, we assume that the ratio of the mutant copy number to the total copy number in the tumour remains the same in the blood plasma, which should be equivalent to the ratio of tumour-derived mutant reads to total tumour-derived reads in the paired plasma WGS.

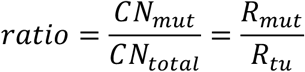

where R_mut_ is the mutant reads at this locus and R_tu_ is the total tumour-derived reads. Note that the total coverage (reads) in the plasma at this locus contains normal-cell-derived wild-type reads and total tumour-derived reads, while the total tumour-derived reads are unknown and contain tumour-derived wild-type and mutant reads. Therefore, the total tumour-derived reads can be calculated as:

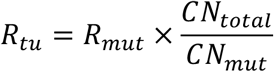

Finally, by aggregating all the n loci with SNVs, plasma TFx can be calculated as:

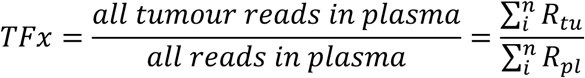

where R_pl_ is the total coverage at each locus.

### Calculation of background plasma TFx

Tumour-derived SNV can be detected by chance in any plasma WGS due to random non-tumour-derived mutations or sequencing artifacts. To account for this noise, the background plasma TFx was calculated. MRDetect was used to search for each patient’s tumour SNV compendium in 32 healthy control plasma WGS. Interestingly, we found that many detected SNVs in control plasma had high VAFs around either 0.5 or 1 (**Supplementary Fig. 2b**). Using the paired buffy coat WGS of the control plasma, we confirmed that these somatic tumour SNVs from a patient happened to be germline single nucleotide polymorphisms (SNPs) in the healthy controls. This would falsely elevate the background TFx, so we removed the variants with depth < 10 reads and VAF ≥ 0.3 from the calculation of background TFx to eliminate the potential germline SNPs. Then, the patient specific TFx detection threshold was determined by the TFx value at 95% z score modeled from background TFx of 32 controls. Finally, the TFx in a patient’s plasma was adjusted by subtracting this threshold. A positive adjusted TFx implies positive detection of ctDNA with 95% specificity. The adjusted TFx was used throughout the downstream analysis in the current study.

#### *In-silico* validation of current plasma TFx algorithm

In-silico plasma WGS samples were generated according to an approach previously described in Zviran et al. 2020^2^. Briefly, paired tumour tissue and buffy coat WGS alignment data from each of 5 PDA patients were downsampled and merged using samtools (version 1.15.1)^59^. Due to the impurity of the tumour sample, the fraction of the reads in the tumour WGS that are actually from the tumour cells (T_TF_) can be described as below:

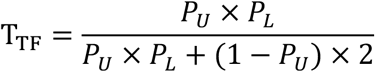

where P_U_ is the purity of the tumour sample and P_L_ is the ploidy of the tumour. Both P_U_ and P_L_ were estimated by CELLULOID.

To downsample the tumour WGS to simulate a plasma WGS with a defined plasma tumour fraction (TFx), a scaling factor (S), is applied:

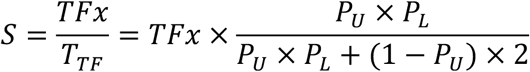

Finally, the downsampling ratios from the tumour and buffy coat WGS can be calculated as below:

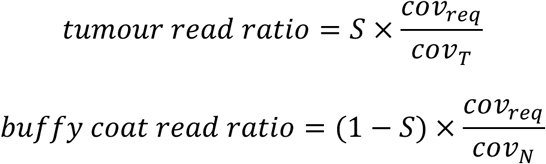

where cov_req_ is the coverage of simulated plasma WGS, cov_T_ is the coverage of the tumour WGS and cov_N_ is the coverage of the buffy coat WGS.

Using the above approach, we simulated plasma WGS data with a range of TFx (1×10^-4^ to 0.5) and sequencing depth (1 to 50x), followed by conducting SNV callings using MRDetect and TFx estimation using our algorithm with different numbers of SNVs input (1,000 to 50,000) (**Supplementary Fig. 2c**). In the in-silico data, we demonstrated that the performance (accuracy and sensitivity) of TFx estimation increased with sequencing depth and the number of SNVs tracked. Even in the low-pass WGS (1x), our approach still had decent performance at TFx of 5×10^-3^. According to the sequencing depth (20 to 60x) and input SNV numbers (∼5,000) of the current PDA cohort, the lowest limit of detection (LOD) of our algorithm is 5×10^-4^ which is similar to the LOD of the tumour fraction formula using the Bernoulli model in Zviran et al. 2020.

### Benchmark of approaches for estimating plasma tumour fraction using real plasma WGS

The tumour fraction values calculated using the current plasma TFx algorithm, TF_ber_ from Zviran et al. 2020 and ichorCNA were compared in the baseline plasma of the current PDA cohort (**Supplementary Fig. 2d**). We computed TF_ber_ without the noise term (µ×R), while instead the TF_ber_ was adjusted by subtracting the TF_ber_ value at 95% z score modeled from background TF_ber_ of 32 controls to keep the normalization step consistent with the current plasma TFx algorithm.

The three approaches were significantly concordant in the general trend (0.73-0.87 for Spearman’s correlation), but some distinct differences were seen. The maximum values in TF_ber_ (∼0.08) were greatly lower than those in the current approach (∼0.7) or ichorCNA (∼0.3). TF_ber_ does not account for heterozygosity and subclonality so it can underestimate the plasma tumour fraction, especially for high tumour fraction samples. Although our plasma TFx approach requires additional data processing for CNVs and VAFs, it considers the wild-type copies from the tumour and adjusts the VAFs of SNVs for their subclonality in the tumour. It can provide an accurate estimation of the real tumour burden (especially for higher levels), so it allows better interpretation for the results when comparing plasma tumour burden under different clinical or tumour molecular conditions. Even though the VAFs in plasma can sometimes be variable in the low tumour fraction settings, we reason that by aggregating thousands of SNV loci, the overall estimation of the final tumor fraction is still accurate. ichorCNA does not require a paired tumour WGS, but has a higher LOD (∼0.03 according to the original publication^58^), which is challenging for analyzing low ctDNA shedders.

### Calculation of short to long ratio in plasma

The genome-wide chromatin openness of the cells of the origin in the plasma was estimated using the DELFI approach which was originally established in Cristiano et al. 2019^3^. Briefly, the function, *delfi*, from FinaleToolkit^60^ was used to calculate the short to long ratio (SLR). The hg38 human genome was divided into non-overlapping 100 kb bins. Reads that fell into the regions in the ENCODE blacklist^61^ (https://www.encodeproject.org/files/ENCFF356LFX/@@download/ENCFF356LFX.bed.gz) were excluded (*--balcklist-file*). The hg38 gap file was downloaded from the UCSC Genome Browser^62^ and provided for *--gap-file* to exclude the regions of centromere, telomere and short arms. The short and long cfDNA fragments were defined as having the lengths between 100-150 bp and 151-220 bp, respectively. GC correction was conducted to account for the bias in the fragment coverage caused by different GC contents across bins. Locally weighted scatterplot smoothing (LOWESS) with a span of ¾ was performed to the data of fragment coverage as a function of the mean fragment GC content in each of the 100 kb bins. The coverage prediction from the LOWESS model was subtracted from the raw coverage to obtain the residuals which were independent of the GC contents of the bins, and the residuals were again normalized by adding back the median of the raw genome-wide coverage. This GC correction was conducted for the short and long fragments separately. To reduce the noise, the data of corrected fragment coverage from 100 kb bins were merged into 5 Mb bins. The SLR in each 5 Mb bin was calculated by the corrected number of short fragments divided by the corrected number of the long fragments in that bin. Finally, the SLR was centered by subtracting the mean of SLRs across all 5 Mb bins for each sample.

### Inference of TFBS accessibility through cfDNA coverage

Transcription factor binding sites (TFBSs) were downloaded from the GTRD database^63^. Following filters were applied: (1) Transcription factors (TFs) had to present in the list identified by Lambert et al. 2018^64^ (Table S1: *‘Is TF?’ == ‘Yes’ & ‘TF assessment’ == ‘Known motif’*); (2) Only the TFBSs on the autosomes were analyzed and only the TFs with more than 10,000 TFBSs on the autosomes were retained; (3) The TFBSs were sorted descendingly by the ‘*peak.count’*, and the top 1,000 TFBSs were retained for each TF. These filters resulted in 555 TFs. The filtered TFBSs were provided to Griffin^65^ to calculate the GC-corrected central coverage at each TFBS with default parameters.

### cfDNA end-motif

A customized python script was created to profile cfDNA fragment end motifs according to Jiang et al. 2020^4^. Briefly, reads on the autosomes were examined, properly paired reads in the primary alignments were retained and the reads that were PCR-duplicated and failed the QC were removed. Fragments with an inferred insert size between 100 and 220 bp were analyzed. The combination of the first 4 nucleotides (4-mer) at the 5’ end of each cfDNA fragment was determined, and all 256 possible combinations of 4-mers were counted for each plasma. The base quality of all the 4 bases in the 4-mer had to be larger than 20 to be considered in the calculation.

### Machine learning classifier for PDA detection using cfDNA fragmentomic features

The machine learning procedure was conducted in Python (version 3.11.0) using scikit-learn (version 1.6.1)^66^. Patients with longitudinal plasma and controls with previous history of cancer were excluded in the training process, which ended up with 151 PDA and 32 control plasma (total n = 183) for building the classifier. A bootstrap sampling strategy was used in training. For each iteration, n samples were randomly selected with replacement for cross-validation (CV), while the samples that were not selected during bootstrapping were used as a held-out test set. SLRs of the 512 non-overlapping 5 Mb genomic bins were used as features, and each feature was z-standardized. To further reduce the feature dimension, Principal Component Analysis (PCA) was used and the top principal components with 80% explained variances were maintained in the model. The standard scaler and PCA were only fitted by the training data in CV, and then the fitted ones were applied to the validation data in CV and the held-out test set to avoid data leakage. A logistic regression with L1 regularization, i.e., the least absolute shrinkage and selection operator (LASSO) regression, was used for modeling. Stratified 5 fold CV was conducted on the data selected by bootstrapping to optimize the hyperparameter, *C* (inverse of regularization strength), in a grid search “*C={0.001, 0.01, 0.1, 1, 10}*”. Area Under the Curve (AUC) was targeted for optimization and “*class_weight = ‘balanced’* ” was selected to account for unbalanced classes of patients and controls. The model with the optimized *C* was evaluated in the held-out test set. This process was repeated for 100 iterations, the value of *C* with the most votes among all iterations was selected for the final model. The final model was retrained on the entire dataset and was further used to predict PDA in the held-out longitudinal plasma.

### Quantification of SVs and detection of SVs in plasma

Evidence for SV calls was directly pulled from the BAM treating each breakpoint separately (**Supplementary Fig. 14**). A minimum of 8 bases needed to be soft clipped and matched the reference at the other breakpoint. Discordant pairs needed to cross the breakpoint in the correct distance or orientation. Events smaller than 2,500 bp were removed as these sites can be difficult to find evidence of discordant pairs. The wildtype copies were calculated as any fragments that crossed the breakpoint site without evidence for the SV. The VAF of each breakpoint was calculated as the number of unique breakpoint supporting fragments divided by the total number of fragments crossing the breakpoint site. This was run on the bulk tumour samples to determine the VAFs of SV breakpoints in the tumour.

The copy number of the breakpoint was identified by comparing the site to the tumour segment copy number (CN) determined by CELLULOID. To do this we identified the CN at the breakpoint site along with the neighboring CN states. SV breakpoints are often associated with a copy number changes thus if a copy number change occurred within 1kp of an SV breakpoint we assumed they were the same event. Soft clipped bases represent the actual SV event and occur on the side of the event (i.e., a deletion with soft clips to the right represents loss of the region to the right). In cases where a CN change and an SV breakpoint occur within 1kb of each other, the true CN state is the CN opposite to the clipped bases (i.e., the CN state prior to the SV event). If a copy number change was not found within 1k of the breakpoint, the CN at the exact breakpoint was used. Event where a copy number could not be determined were excluded.

Using the tumour as a guidance, the SV breakpoints were examined in the paired plasma WGS BAM using the same criteria to search for evidence of SV in the plasma.

### Identification of clonal and subclonal SNVs and SVs in tumours

To classify SNVs/SVs as being clonal or subclonal in bulk tumour WGS, we first calculated the mutant counts/copies for all SNVs and SVs as described in our previous work^22^. Briefly, the mutant copies for each alternation were calculated using the formula (1) and (2) mentioned above. Next, we ran mclust v6.1^67^ on the distribution of mutant copies to identify the peaks (**Supplementary Fig. 11 and 15a**). For SNVs, this was run separately for each patient. Due to the low number of SVs found in each individual patient, SVs were merged across the cohort for this step. To optimize the ability to detect clusters, we focused on areas of the distribution where the vast majority of variants would fall (i.e., those with mutant copies less than 4.7). Variants with total depth of less than 30 were treated as noise in the algorithm. 18 fitted models were generated and each model was ranked based on Bayesian Information Criterion (BIC), Integrated Complete Likelihood (ICL) and median uncertainty with 1 being the best model. For each model, the ranks from the 3 metrics were summed and the model with the highest summed rank was considered the best overall model. In the case of ties, the data was manually reviewed to select the best model. For SNVs the subclonal clusters were considered clusters with the median peak less than 0.8. For SVs the subclonal clusters were considered clusters with median peak less than 0.5.

### Validation of detection of subclonal SNVs in plasma

*In-silico* validation of tumour subclone detection was conducted to confirm the over- and under-representation of subclonal SNVs in plasma with low and high TFx, respectively. Three different approaches were used for this purpose.

#### Sampling mutant reads

For a tumour sample with N SNVs successfully assigned as either clonal or subclonal and the total number of mutant reads, J, to support these SNVs, mutant reads were randomly sampled without replacement until the designated number of mutations, M, are found. The number of randomly sampled reads, j, was between M and J (M ≤ j ≤ J) based on uniform distribution. An SNV locus was determined as detected if it had at least 1 mutant read. In each experiment, the proportion of detected subclonal mutations was calculated as the number of detected subclonal SNVs divided by the total number of detected SNVs. For the same M, the same experiments were conducted 200 times to obtain confidence interval. Multiple target numbers of mutations were used to see the trend, for example, every 500 mutations. See the result in **Supplementary Fig. 13a.**

#### Sampling loci

For each patient, the number of subclonal SNV loci detected in plasma (X) was simulated using a hypergeometric distribution:

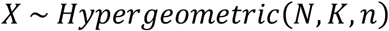

where N is the total number of SNVs in the paired tumor that were successfully assigned as either clonal or subclonal, K is the number of subclonal SNVs in the tumor, and n is the total number of SNVs detected in plasma. The number of subclonal SNVs detected in plasma was randomly generated 1,000 times. These numbers are then divided by n to obtain the simulated subclonal detection rates in plasma. According to the 1,000 simulated detection rates, 95% and 5% quantiles were calculated. The observed (actual) subclone detection rate in plasma was then compared to the quantiles to define whether subclonal SNVs were statistically enriched or depleted in plasma. See the result in **Supplementary Fig. 13b.**

#### MRDetect is not biased towards subclonal mutations

To validation that MRDetect is not biased towards detecting subclonal mutations in plasma, we examined the distribution of supporting mutant reads in plasma. For each SNV, the number of supporting mutant reads were identified. The proportion of x supporting reads were calculated across three TFx groups and separated by clonal and subclonal mutations. The Wilcoxon signed-rank test was conducted between clonal and subclonal groups. See **Supplementary Fig. 13c.**

### Clonal analysis in longitudinal plasma samples

To explore the longitudinal samples in each patient, paired tumours were used as a guidance to search for SNVs in plasma by MRDetect, as described above. This data was combined to provide a complete picture of the tissue-specific mutations found in the plasma. To track bulk tumour evolution, copy number variations were compared across all bulk samples from the same patient. To identify which tumour seeded the others, we focused on sites with loss of heterozygosity (LOH) in one tumour compared to the other sites. If a tumour had a LOH, it could not seed another tumour that missed this LOH event in the same site. Using this logic, we tracked the seeding events (an example in **Supplementary Fig. 19c**). In addition, bulk mutations could be tracked as early (present in the original pancreas tumour) and late (found only in the metastasis). For establishing clones and phylogenetic lineage in PCSI0681, pyclone-vi ^68^ was run.

### Survival and other statistical analysis

All the data analyses were conducted in either R or Python. The Cox Proportional-Hazards model and the Kaplan-Meier estimator were used for survival analysis, which was conducted by survival (version 3.8.3)^69^ and survminer (version 0.5.0)^70^ R packages. All the heatmaps were generated by ComplexHeatmap (version 2.6.2)^71^.

## Code availability

Our novel algorithm for plasma TFx estimation, other custom code and scripts can be accessed at: https://github.com/robinycfang/plasma_WGS_PDA_pub.

## Acknowledgements

We are deeply grateful to the patients and their families for their participation in this study. We would like to thank the members of the Notta Laboratory and the PanCuRx program at the OICR (including Dr. Lincoln Stein) for their valuable discussions, technical input, and feedback on the manuscript. We acknowledge the support of the Princess Margaret Cancer Biobank, the COMPASS trial study team, the Pathology Research Program Laboratory at UHN, and the clinical staff at the Wallace McCain Centre (Pancreatic Cancer Clinic) for their contributions to sample acquisition, processing, and consenting patients. This work was supported by the PanCuRx Translational Research Initiative funded by the Government of Ontario through OICR, Wallace McCain Centre, Princess Margaret Cancer Foundation, Canadian Cancer Society Research Institute, Canadian Institutes of Health Research, Ontario Ministry of Economic Development, and Pancreatic Cancer Canada. We are also thankful for philanthropic support from the Canadian Friends of the Hebrew University (A. U. Soyka). F.N. is supported by the Gattuso-Slaight Personalized Cancer Medicine Fund, the Ontario Institute for Cancer Research, the Canadian Institutes of Health Research (grant no. 451067), and the Canada Research Chairs Program. Y.F. obtained a scholarship from Marathon of Hope Cancer Centres Network.

## Contributions

Y.F., M.C. and F.N. conceptualized the project. K.N., D.B., S.R., A.B., S.H., A.D., Y.W., S.H., R.G., E.S.T., G.Z., G.M.O, J.J.K., S.G. and F.N. collected the patient samples. T.H., C.G., D.B., S.R., A.B., S.H., A.D., M.A.H., K.N. and Y.F. obtained and compiled the patients’ clinical records. Y.F., M.C., A.Z., G.H.J., S.C. and J.M.W. managed and processed the raw sequencing data. Y.F. and M.C. performed most of the data analysis and interpreted with F.N.. A.Z., G.H.J. and S.W.K.N. provided additional bioinformatic analysis. E.F., A.E., M.M. and B.T.G. created and annotated histology slides. Y.F., M.C. and F.N. wrote the manuscript, with feedback from A.Z., E.F. and S.G..

---

**Supplementary** Fig. 1 | Molecular profiling from the paired tumour tissue WGS and RNA-seq in the current pancreatic cancer cohort. **a**-**c**, The numbers of SNVs, small insertions and deletions (INDELs) and structural variations (SVs) in the tumours. **d**-**g**, The distribution of mutation types in the common pancreatic cancer drivers. In KRAS, the balanced type refers to the equal numbers of mutant and wild-type (WT) alleles, while the minor and major imbalanced types refer to favoring the mutant alleles over the WT ones slightly and considerably, respectively^25^. In other drivers, “biallelic; mut” refers to situations where either both alleles are mutant or one mutant allele is present and WT allele is lost due to loss of heterozygosity (LOH), while “biallelic; homdel” refers to a homozygous deletion of both alleles. **h**-**i**, The distribution of homologous recombination deficiency (HRD) and mismatch repair deficiency (MMR) in the tumours. **j**, The cohort-wise frequencies of copy number gain and loss in 5 Mb bins across the genome in the tumours. **k**-**i**, The distribution of ploidy types (with a cut-off of 2.3 for total tumour copies) and transcriptional subtypes (according to Moffitt et al. 2015^13^) in the tumours. All the associations between molecular features and tumour tissue sites were evaluated using the Fisher’s exact test.

**Supplementary** Fig. 2 | Development and validation of the algorithm for plasma tumour fraction (TFx) estimation. **a**, The schematics for tumour-guided mutation calling and TFx estimation in plasma. MRDetect was used to search plasma WGS for SNVs identified from the paired tumour WGS. The information of detected SNVs in plasma, along with SNVs, copy number variations (CNVs) and tumour tissue cellularity, was integrated in a formula to estimate plasma TFx. **b**, Two examples for detection of SNVs from 2 tumour tissues in 32 non-tumour control plasma samples. The y axis indicates the variant allele frequencies (VAFs) of detected SNVs in controls. Many detected SNVs showed markedly high VAFs, implying that these SNVs that were somatic mutations in tumours might accidentally be germline single nucleotide polymorphisms (SNPs) in the controls. SNVs with a VAF > 0.3 (indicated by a red horizontal dashed line) were defined as potentially germline in controls and were removed from calculation of the background TFx in controls to avoid falsely high background levels due to inclusion of germline SNPs. **c**, Performance evaluation of the algorithm to estimate plasma TFx using *in-silico* plasma data. Plasma WGS data with different levels of TFx and sequencing coverage were simulated by mixing reads from a tumour WGS and its paired buffy coat WGS. Different numbers of input SNVs were also tested for plasma TFx. In total, 5 donors were used in this simulation, and the dots and error bars represent medians and quantiles (25% and 75%) of the estimated plasma TFx. The data highlighted by the green box indicate the range of lowest limit of detection (5×10^-4^ to 1×10^-3^) given the SNV numbers and sequencing coverages in the current pancreatic cancer cohort. **d**, The benchmarking of different TFx approaches (plasma TFx, the tumour-guided formula in the current study; TFx, Bernoulli, another tumour-guided formula using the Bernoulli model in Zviran et al. 2020^2^; ichor TFx, tumour-agnostic ichorCNA). For each approach, a background TFx was calculated as the TFx at 95% (one-sided, upper) z score calculated from 32 control plasma, and this background TFx was subtracted from the TFx value of each patient to adjust for the background noise. The adjusted TFx that were negative were set to 0 and they indicated no detection of ctDNA. The Spearman correlations were used in this analysis.

**Supplementary** Fig. 3 | **a**, Detection of SNVs in plasma using conventional mutation callers. Mutect2 and Strelka2 were conducted to identify somatic SNVs using plasma and paired normal WGS. The SNVs identified in plasma by both callers were further intersected with the SNVs from the paired tumour WGS. **b**, Background cumulative VAF of SNVs detected in controls. For each patient, using the list of tumour SNVs, MRDetect was conducted to identify background SNV detection in 32 non-tumour control plasma samples, followed by calculating cumulative VAFs of the detected SNVs in controls, and comparing the values between different groups of SNV detection rate (according to the tested paired tumour and patient plasma). **c**, Distribution of individual plasma VAFs of detected SNVs from controls, stratified by detection rate.

**Supplementary** Fig. 4 **| a**, ctDNA positivity in treatment-naïve plasma across clinical stages. Samples with the adjusted plasma TFx (background TFx from 32 controls subtracted) large than 0 is defined as TFx positive. **b**, Pre-resection plasma TFx in Stage I and II patients. Pre-resection plasma TFx (**c**) and tumour size measured from CT scans (**d**) for Stage II patients with negative (N0) and positive (N1) lymph nodes. **e**, Pre- and post-resection plasma TFx (unpaired) for early-stage (I and II) patients. **f**, The prognostic values of post-resection plasma TFx in resectable cases. The binary cut-off (low or high TFx level) for optimal prognosis was determined using the Univariate Cox analysis (Supplementary Table 6). The optimal cut-off was then used to stratify the patients and the Kaplan-Meier analysis was used in the survival curves.

**Supplementary** Fig. 5 | Clinical associations of serum CA19-9. **a**, Treatment-naïve CA19-9 across clinical stages. The Wilcoxon signed-rank test was conducted. **b**, Treatment-naïve CA19-9 positivity (> 37 U/mL) across clinical stages. **c**, Pre- and post-resection CA19-9 (paired) for resectable cases (Stage I and II). **d**, The prognostic values of post-resection CA19-9 in resectable cases. The binary cut-off (low or high CA19-9 level) for optimal prognosis was determined using the Univariate Cox analysis (Supplementary Table 6). The optimal cut-off was then used to stratify the patients and the Kaplan-Meier analysis was used in the survival curves.

**Supplementary** Fig. 6 | The Spearman correlations (shown as the numeric values) between treatment-naïve blood biomarkers and tumour sizes measured from CT scans in resectable (**a**), locally advanced (**b**) and metastatic (**c**) cases. In metastatic cases, the sum of mets size is the sum of any measurable lesions.

**Supplementary** Fig. 7 | The correlations between treatment-naïve plasma TFx and serum CA19-9 and tumour sizes measured from CT scans across clinical stages. The Spearman correlations were used. The gray and red dashed lines indicate the positivity thresholds for plasma TFx (0%) and CA19-9 (37 U/mL), respectively. The sum diameter in Stage IV patients indicates the sum diameter of primary tumour and any measurable metastatic lesions.

**Supplementary** Fig. 8 | The plasma TFx levels under different tumour molecular conditions in metastatic patients. The Waddell, Moffitt, Ge, Collisson, Bailey subtypes refers to classification schemes in Waddell et al. 2015^9^, Moffitt et al. 2015^13^, Ge et al. 2026^12^ and Collisson et al. 2019^57^, respectively. The Wilcoxon signed-rank test and the Kruskal-Wallis test were used for comparisons in 2 and 3 groups respectively.

**Supplementary** Fig. 9 | Tumour molecular mechanisms to impact ctDNA shedding in metastatic pancreatic cancer. **a**, Top 10 enriched Hallmark gene sets in polyploidy tumours identified by GSEA. **b**, The Spearman correlations between plasma TFx and immune repertoires in the tumour tissues in metastatic cases. Repertoire abundance was estimated as the number of detected clonotypes (TRA, TRB, TRG, TRD for TCR and IGH, IGK and IGL for BCR), as identified by MiXCR.

**Supplementary** Fig. 10 | cfDNA fragmentomic features in the pancreatic cancer. **a**, The distribution of cfDNA fragment length in plasma of non-tumour controls and pancreatic cancer patients. The curves represent the median values from different groups. **b**, Evaluation of the batch effect (plasma from either Biobank or OPCS) on 3 sets of fragmentomic features (bin-wise SLRs, central coverage at transcription factor binding sites (TFBSs) using Griffin and 4-mer end-motifs). **c**, The volcano plot for central coverage at TFBSs of 555 transcription factors profiled by Griffin. The log2FC was calculated by the log2 of central coverages in patients over those in controls. DOWN, NS and UP indicate that the central coverage at TFBSs was reduced, not different, and increased in patient plasma, compared to control plasma. **d**, The Receiver Operating Characteristic (ROC) curve for the performance of a LASSO regression trained with 512 bin-wise SLRs to classify pancreatic cancer patients from controls. The area under the curve (AUC) was calculated on the held-out test set from each of 100 bootstrap samplings. The model was also tested on the downsampled coverages down to 0.1x. **e**, The sensitivities of the model on the held-out test set at 80% specificity, under different clinical stages and plasma TFx.

**Supplementary** Fig. 11 | Examples of SNV profiles identifying 0, 1 or 2 subclones in diploid and tetraploid tumours.

**Supplementary** Fig. 12 | **a**, Percent of subclonal plasma SNVs across TFx groups within subclone size groups. **b**, Percent of subclonal plasma SNVs across TFx groups with different number of subclones.

**Supplementary** Fig. 13 | Validation of detection of subclonal SNVs in plasma. **a**, Sampling mutant reads. Proportion of subclonal mutations identified when BAM was downsampled to x number of mutations. **b**, Sampling mutation loci. i) All the mutation loci were randomly sampled 1,000 times, followed by calculation of subclonal mutation detection rate to form the background. The grey dot and error bars indicate median and quantiles (5% and 95%) of the background. ii) The summary of statistics for subclone representation in different TFx groups. The observed subclone detection rate from the patient plasma was compared with the quantiles of the background. The observed values larger than 95% and smaller than 5% quantiles of the background were defined as enriched and under representative, respectively; otherwise defined as not significant. **c**, The number of mutant reads supporting a variant separated by TFx group. The Wilcox test was performed.

**Supplementary** Fig. 14 | Screenshots from IGV demonstrating clonal and subclonal structural variant breakpoints.

**Supplementary** Fig. 15 | **a**, Distribution of SV breakpoint mutant copies with the underlying distributions displayed. **b**, Proportion of SV breakpoints across the identified clusters between TFx groups.

**Supplementary** Fig. 16 | Additional information for PCSI0633. **a**, Mutant copies in PCSI0633 pancreas and liver samples. 81% of subclonal pancreas mutations found in the liver are clonal. **b**, Celluloid plots for the pancreas and the liver in PCSI0633. Arrows showing the copy state change between the tumours. **c**, Proportion of mutations identified in plasma samples.

**Supplementary** Fig. 17 | **a**, PCSI0612 profile. i) Clinical timeline. The green segment on the timeline indicates periods when the patient was on chemo- or radiotherapies. ii) Detection rate of tissue-specific SNVs (# detected in plasma / total # in tissue). The background detection rate was calculated by searching tissue-specific SNVs in plasma of 32 healthy controls. The observed patient detection rate exceeding the 95% confidence interval of the background distribution (z > 1.645) was considered significant. iii) Proportion of clonal and subclonal mutations of pancreas specific mutations in plasma. **b**, PCSI1056 profile. i) Clinical timeline. ii) Plasma TFx (%) change over time. iii) Detection rate of tissue-specific SNVs. **c**, PCSI1056 profile. i) Clinical timeline. ii) Plasma TFx (%) change over time. iii) Detection rate of tissue-specific SNVs.

**Supplementary** Fig. 18 | a. PCSI0803 profile. i) Clinical timeline. The green segment on the timeline indicates periods when the patient was on chemo- or radiotherapies. ii) Plasma TFx (%) change over time. iii) Proportion of clonal and subclonal mutations detected in plasma. **b**, PCSI0666 profile. i) Clinical timeline. ii) Plasma TFx (%) change over time. iii) Screenshot of SV breakpoint found in bulk liver sample and plasma1.

**Supplementary** Fig. 19 | Additional information for PCSI0652. **a**, Clonal composition across the clusters identified by pyclone. **b**, Venn diagram of SNVs found in the pancreas, lung and liver of PCSI0652. **c**, The copy number state and the allele frequencies of chr22 in the pancreas, lung and liver.

**Supplementary** Fig. 20 **|** PCSI0473 profile. **a**, Clinical timeline. The green segment on the timeline indicates periods when the patient was on chemo- or radiotherapies. **b**, Detection rate of tissue-specific SNVs (# detected in plasma / total # in tissue). The background detection rate was calculated by searching tissue-specific SNVs in plasma of 32 healthy controls. The observed patient detection rate exceeding the 95% confidence interval of the background distribution (z > 1.645) was considered significant. **c**, Distribution of pancreas specific mutations identified in the plasma samples.

**Supplementary** Fig. 21 | Screenshot of the region of *BRCA2* in PCSI0681. Highlighted are the germline mutation found across all tissues and the reversion mutation found only in the peritoneum.

