## Supplementary Figs. 1-21 for "The subclonal footprint of pervasive early dissemination in pancreatic cancer"

### Supplementary Figure 1

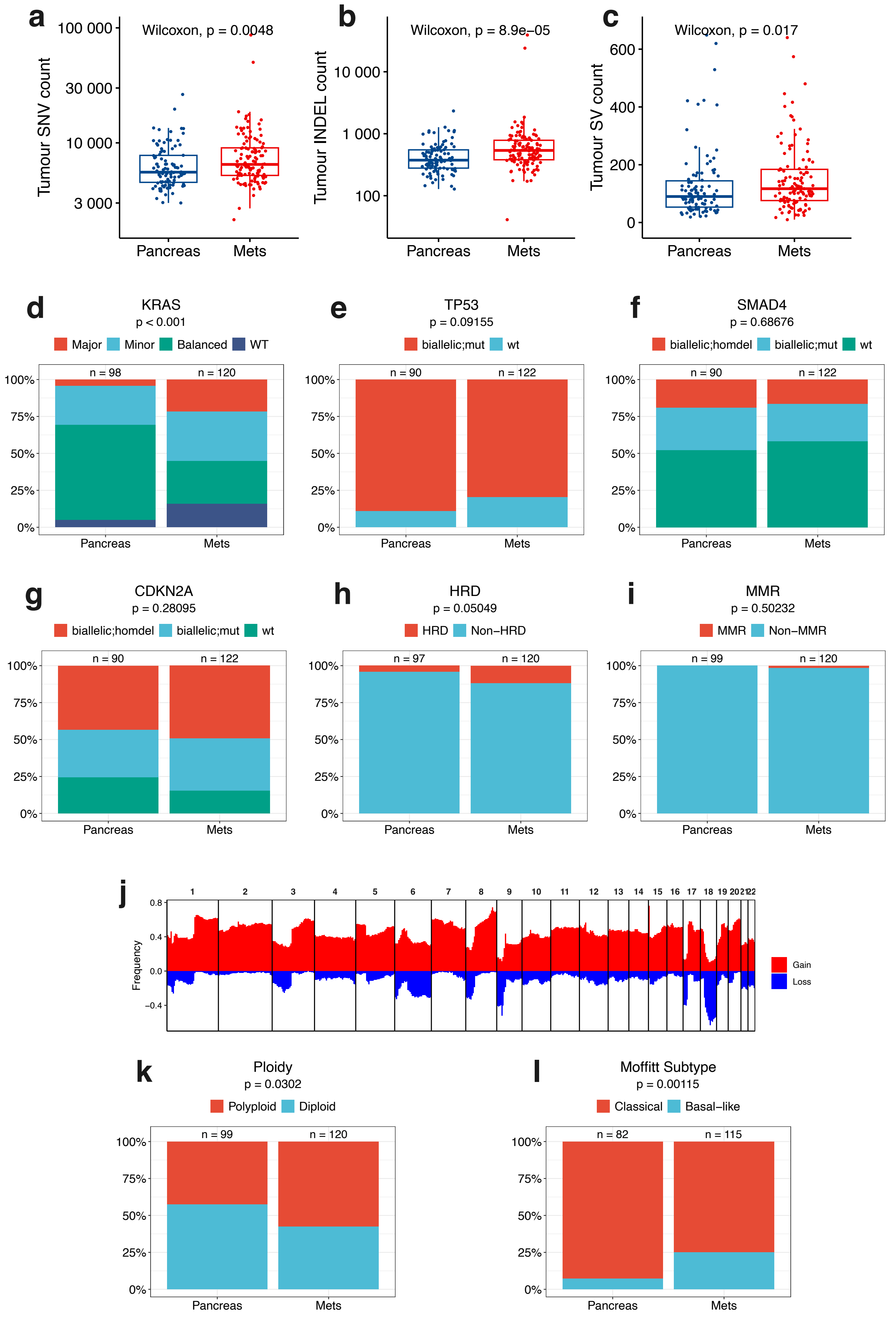

### Supplementary Figure 2

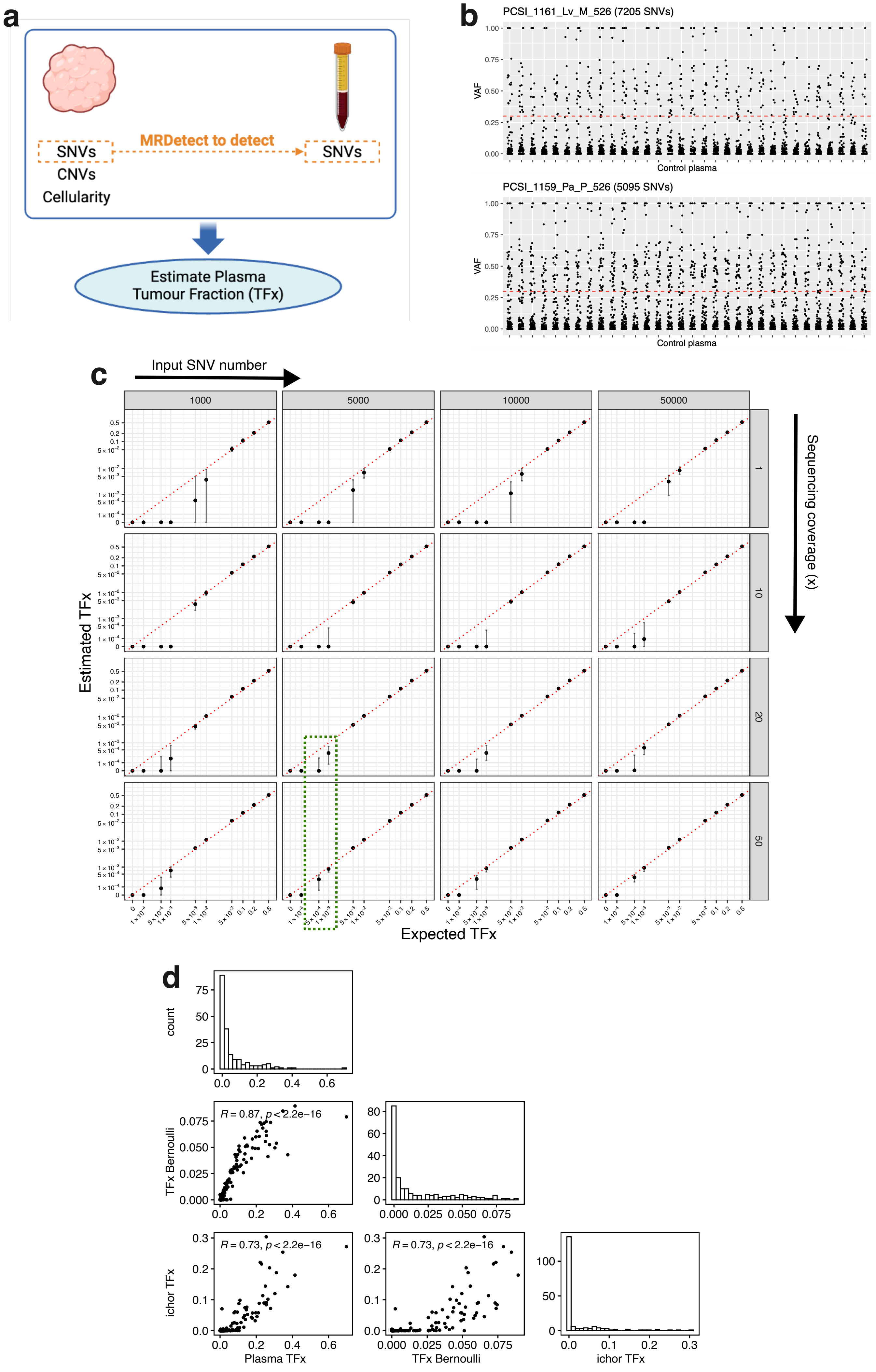

### Supplementary Figure 3

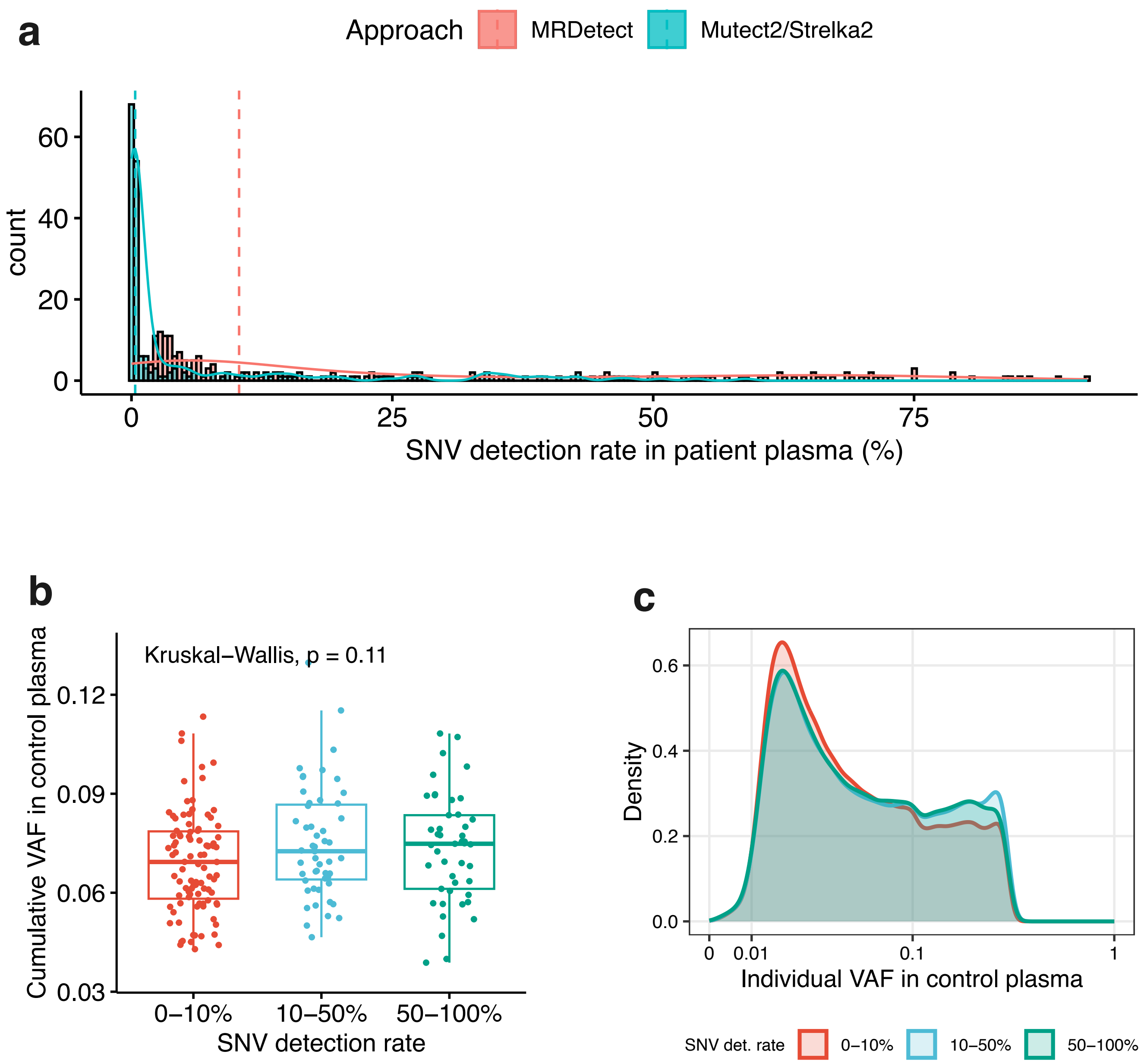

### Supplementary Figure 4

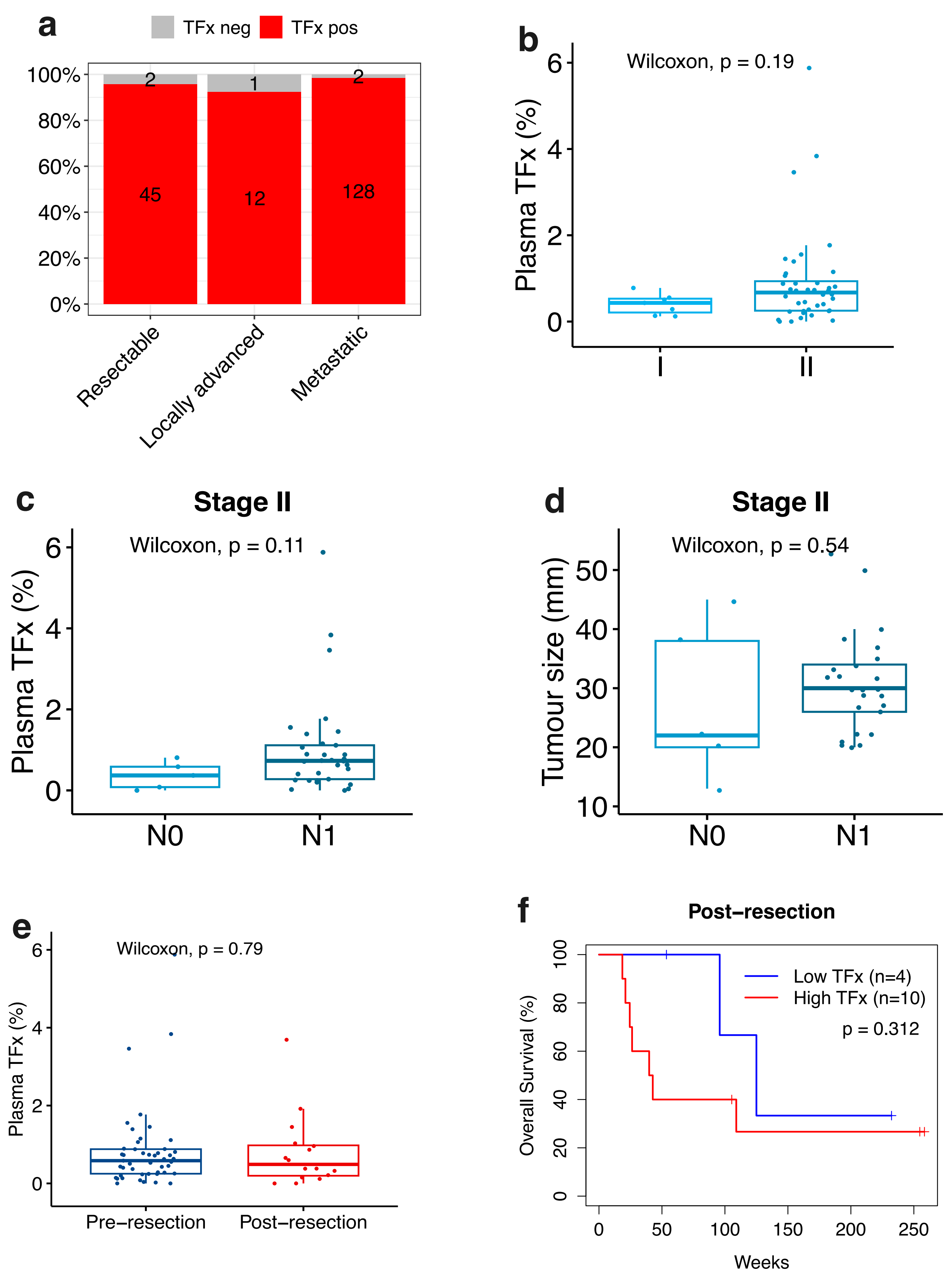

### Supplementary Figure 5

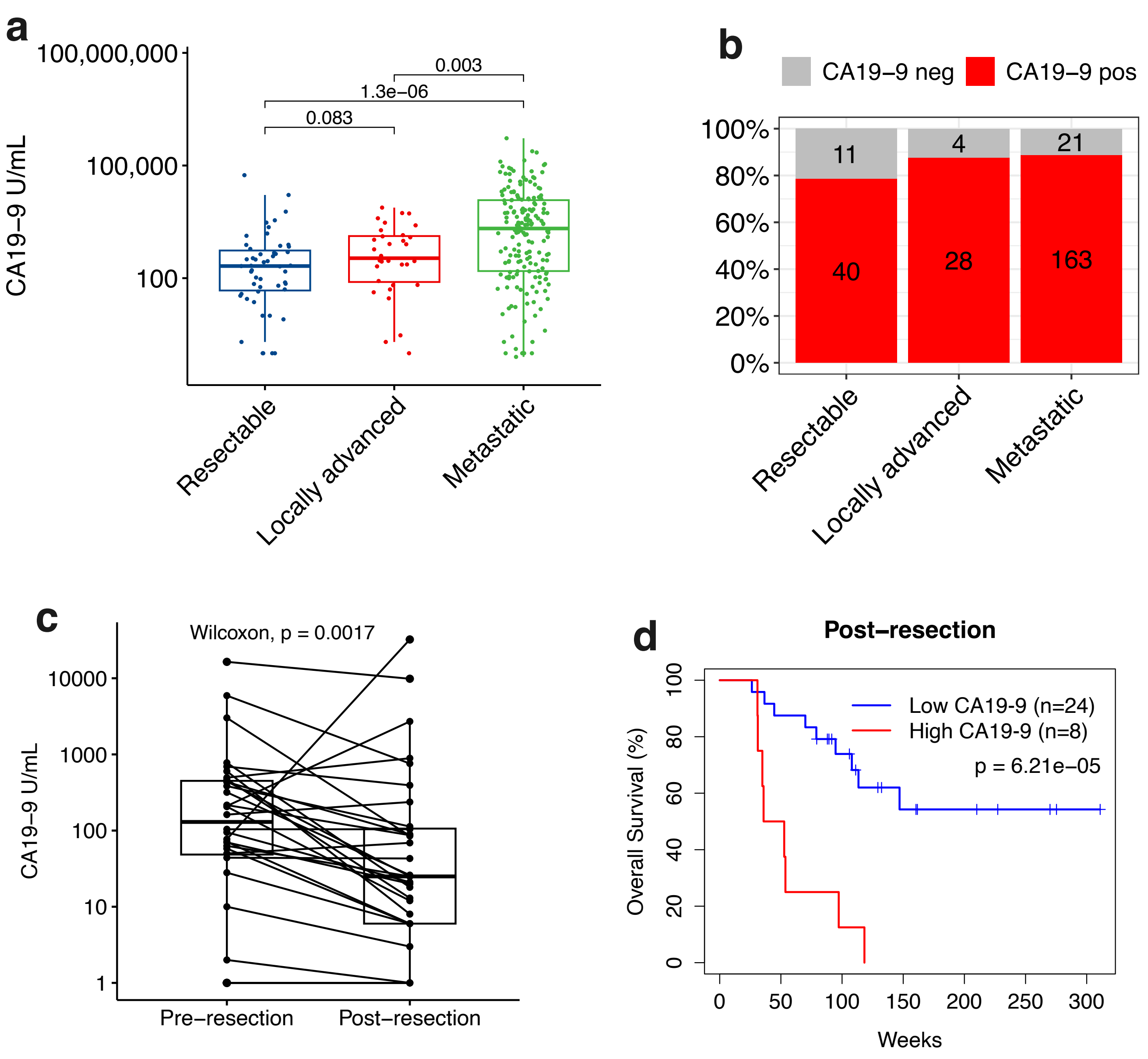

### Supplementary Figure 6

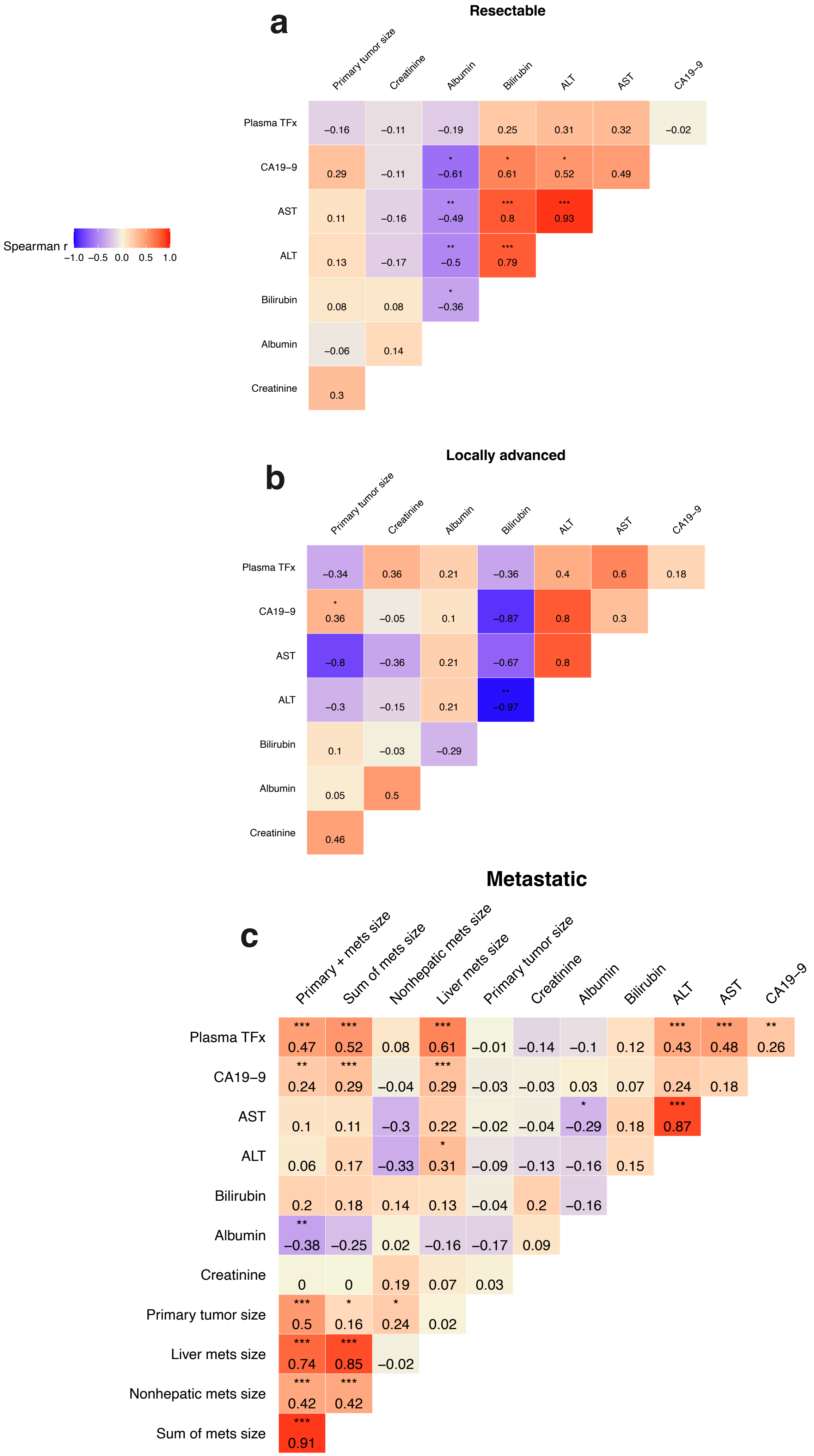

### Supplementary Figure 7

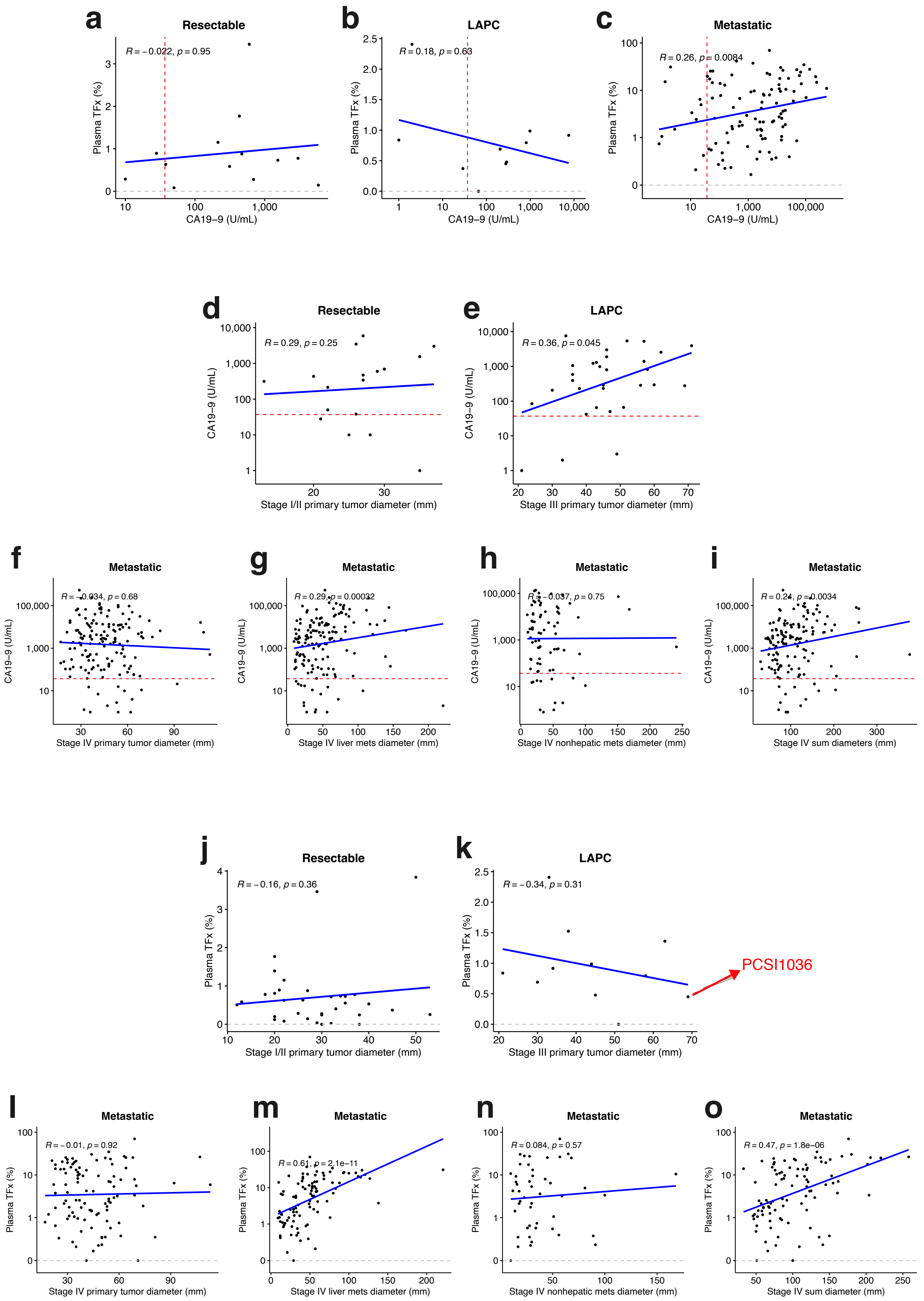

### Supplementary Figure 8

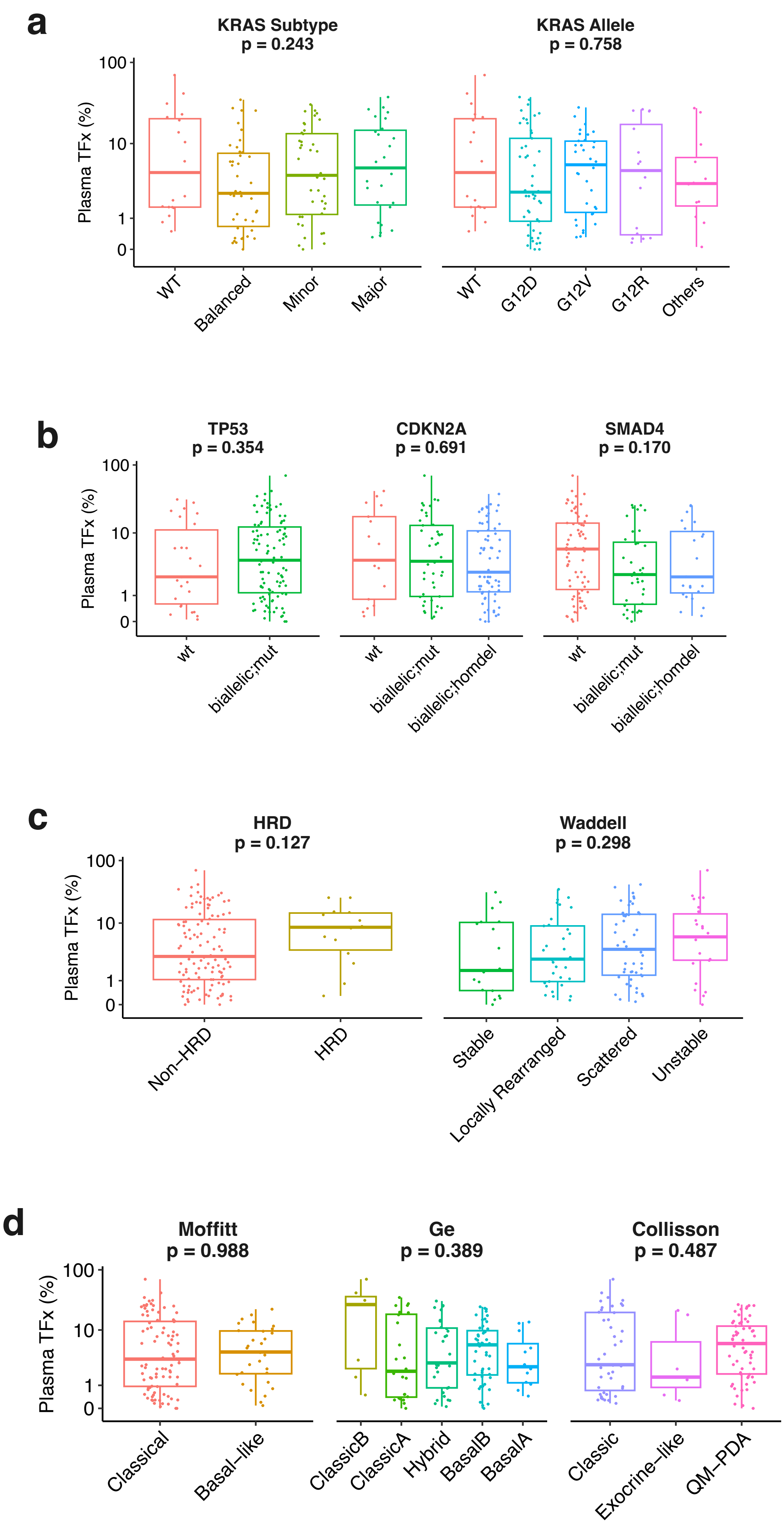

### Supplementary Figure 9

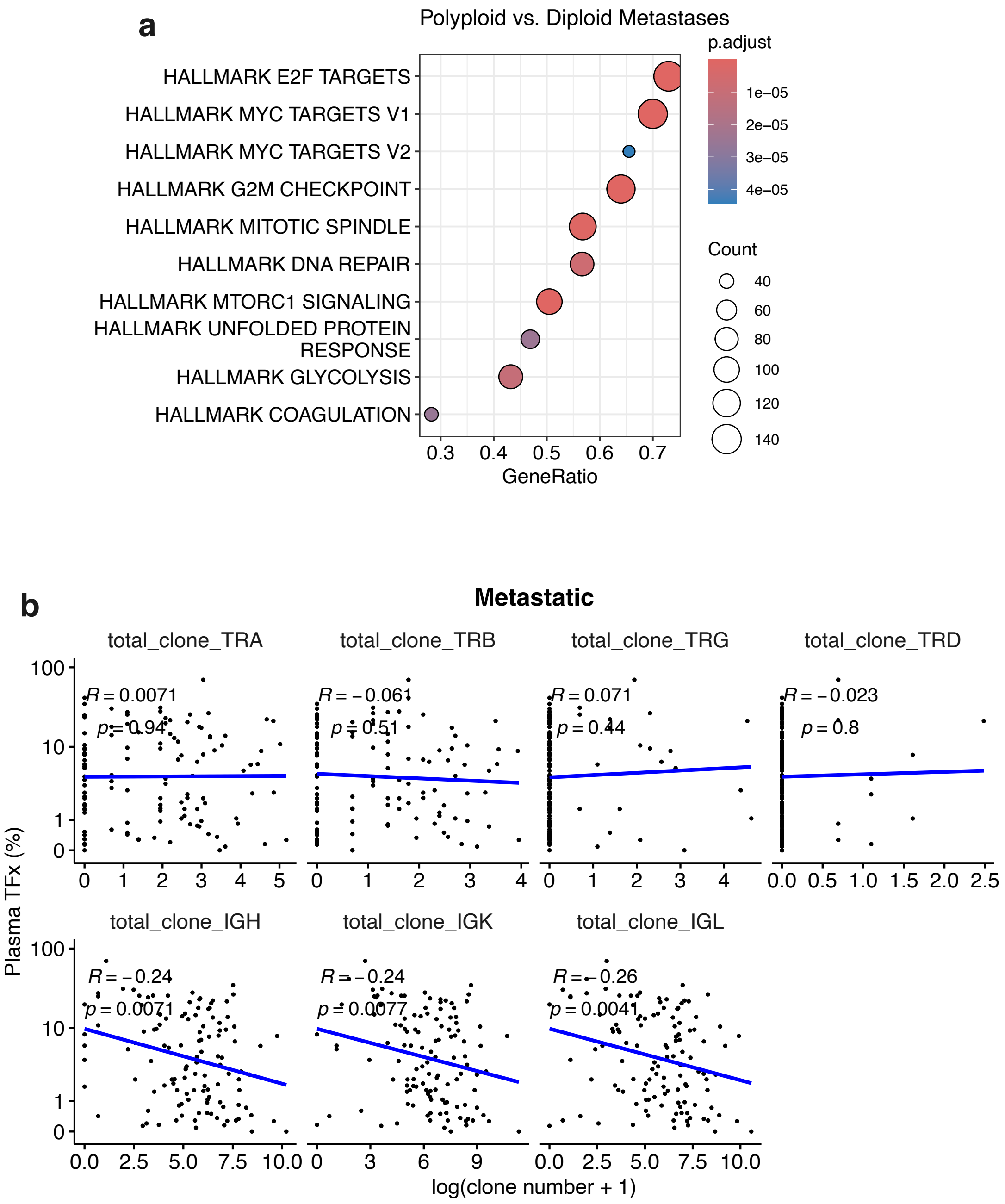

### Supplementary Figure 10

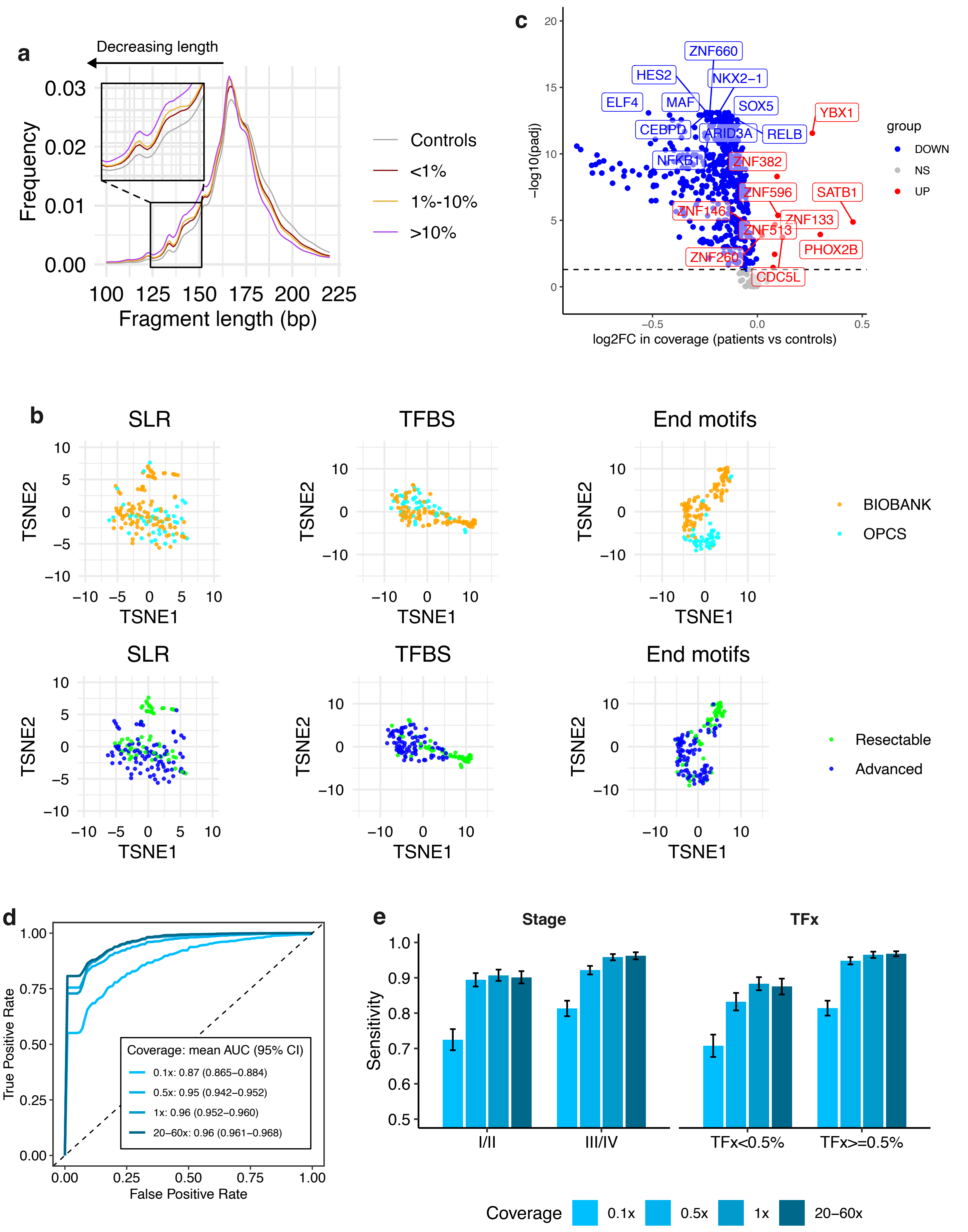

### Supplementary Figure 11

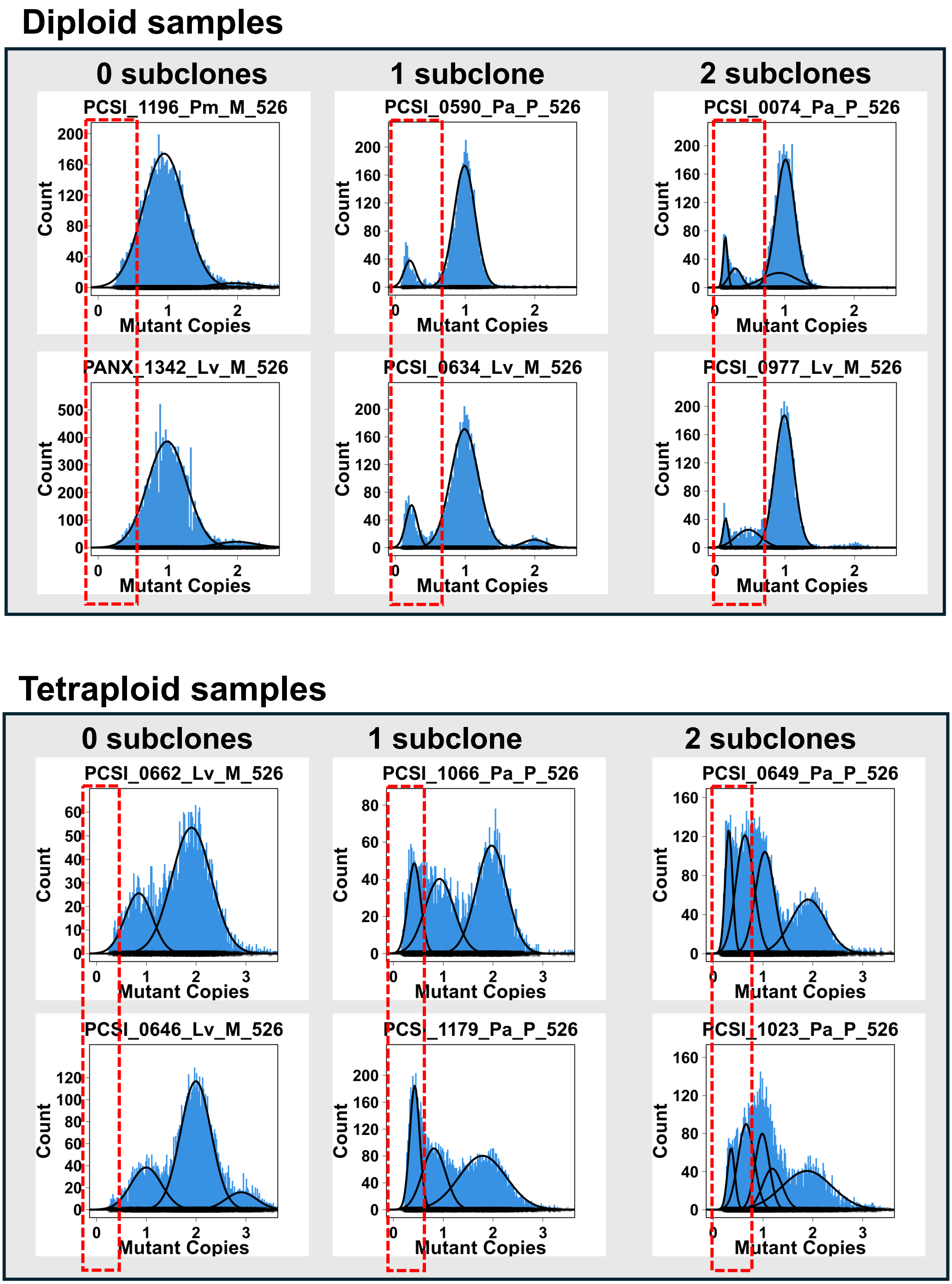

### Supplementary Figure 12

**a**

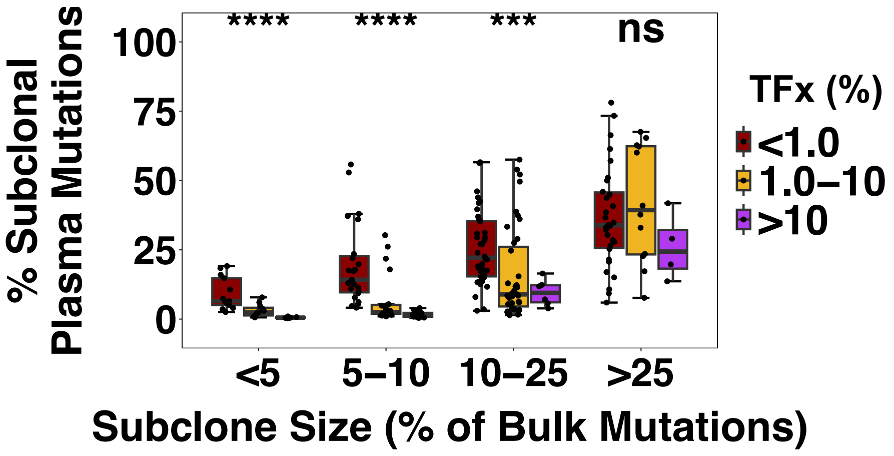


**b**

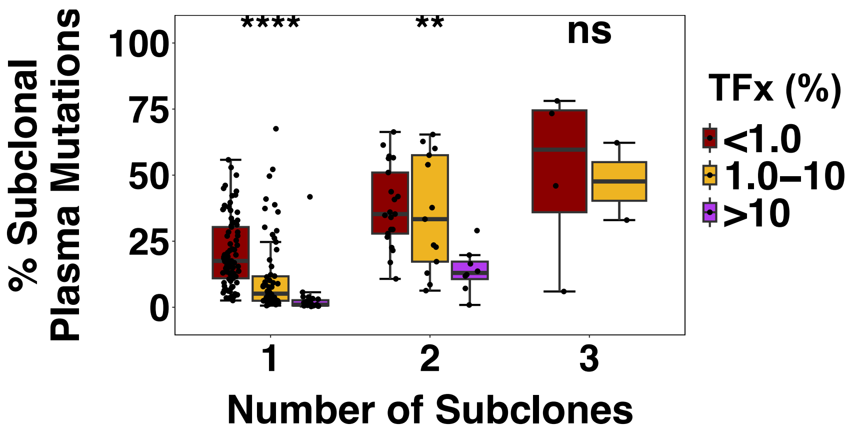

### Supplementary Figure 13

**a**

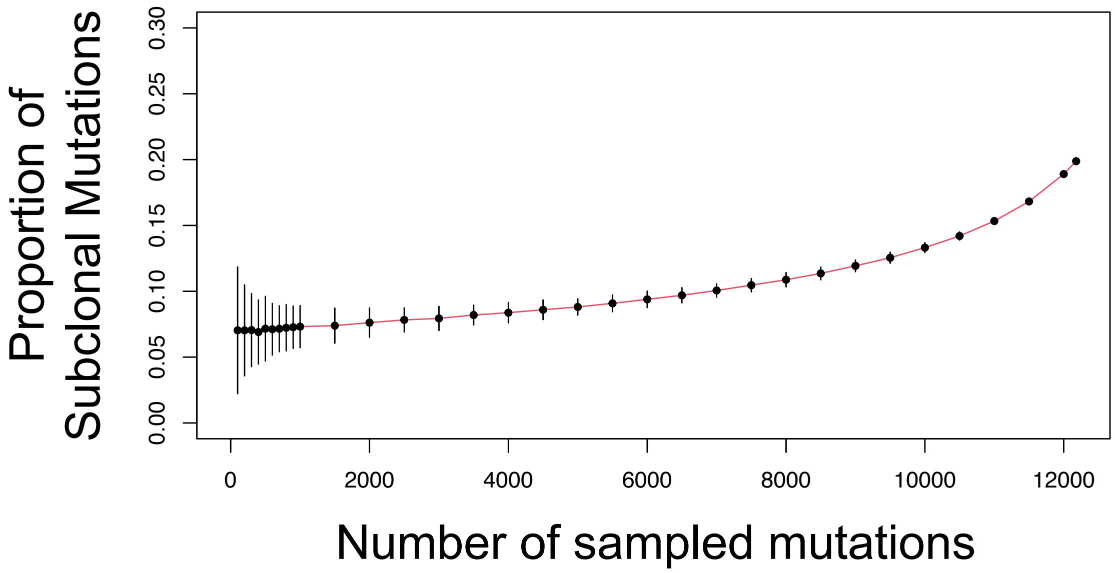


**b**

**i.**

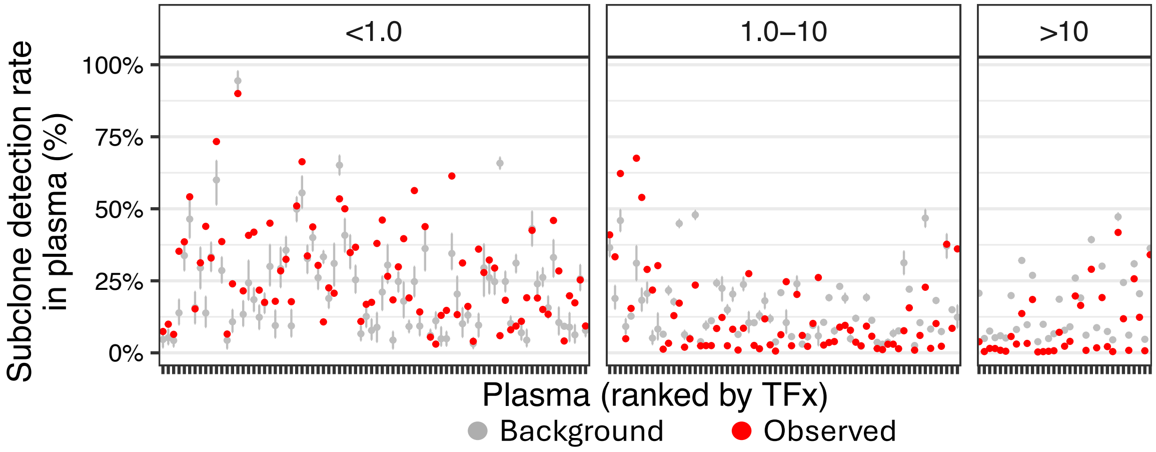


**ii.**

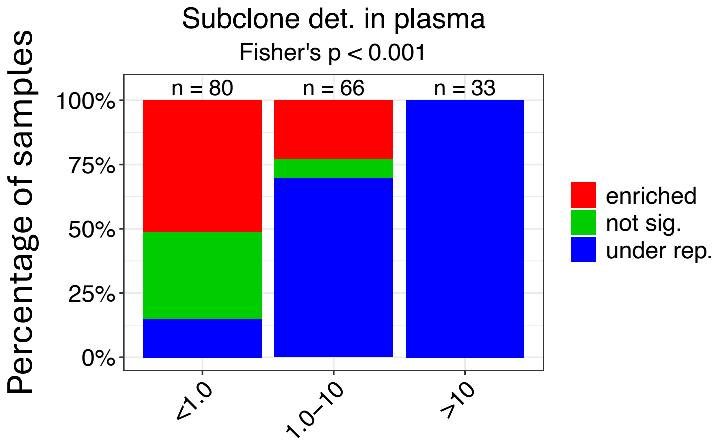

### Supplementary Figure 13 cont'd

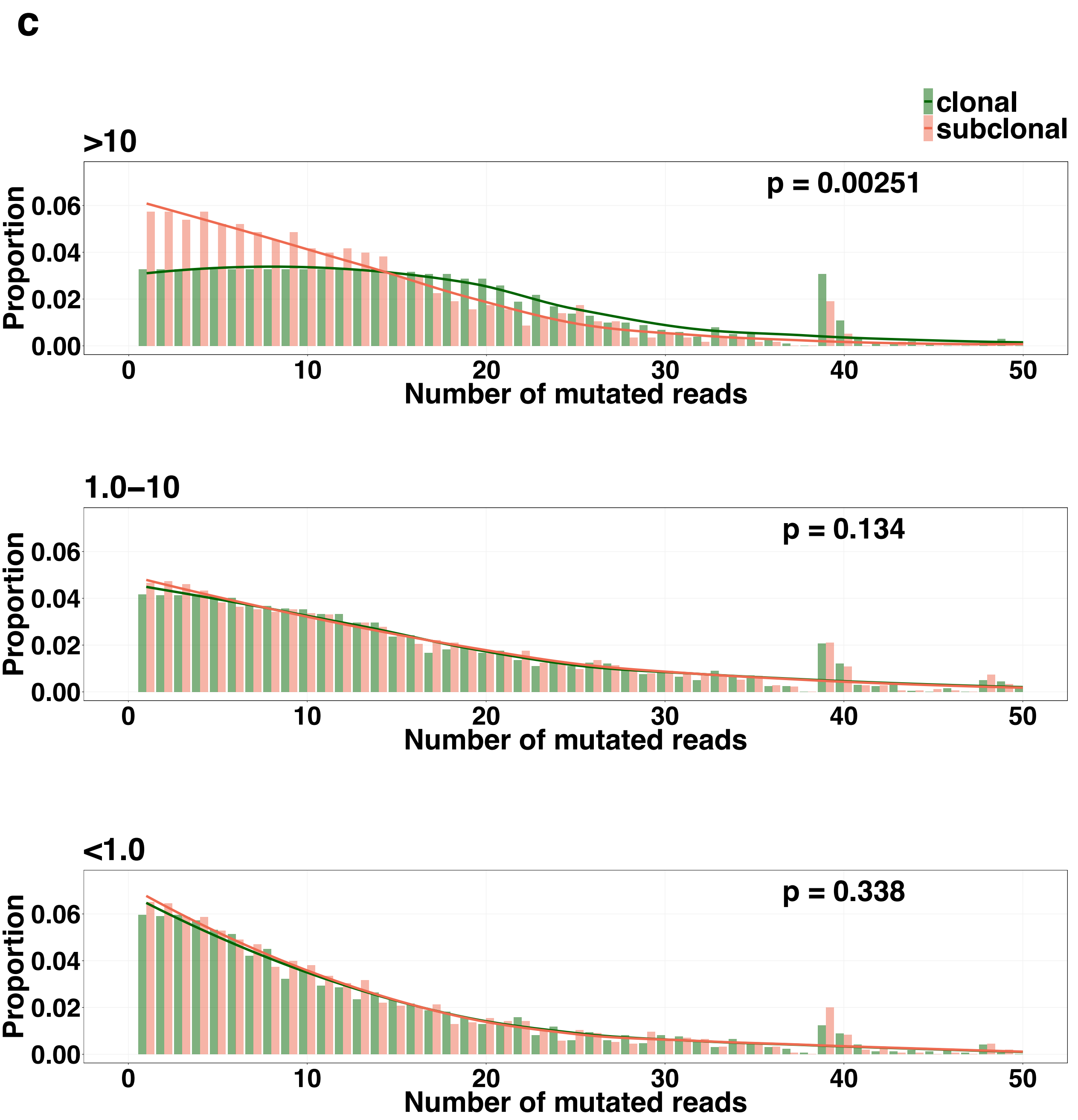

### Supplementary Figure 14

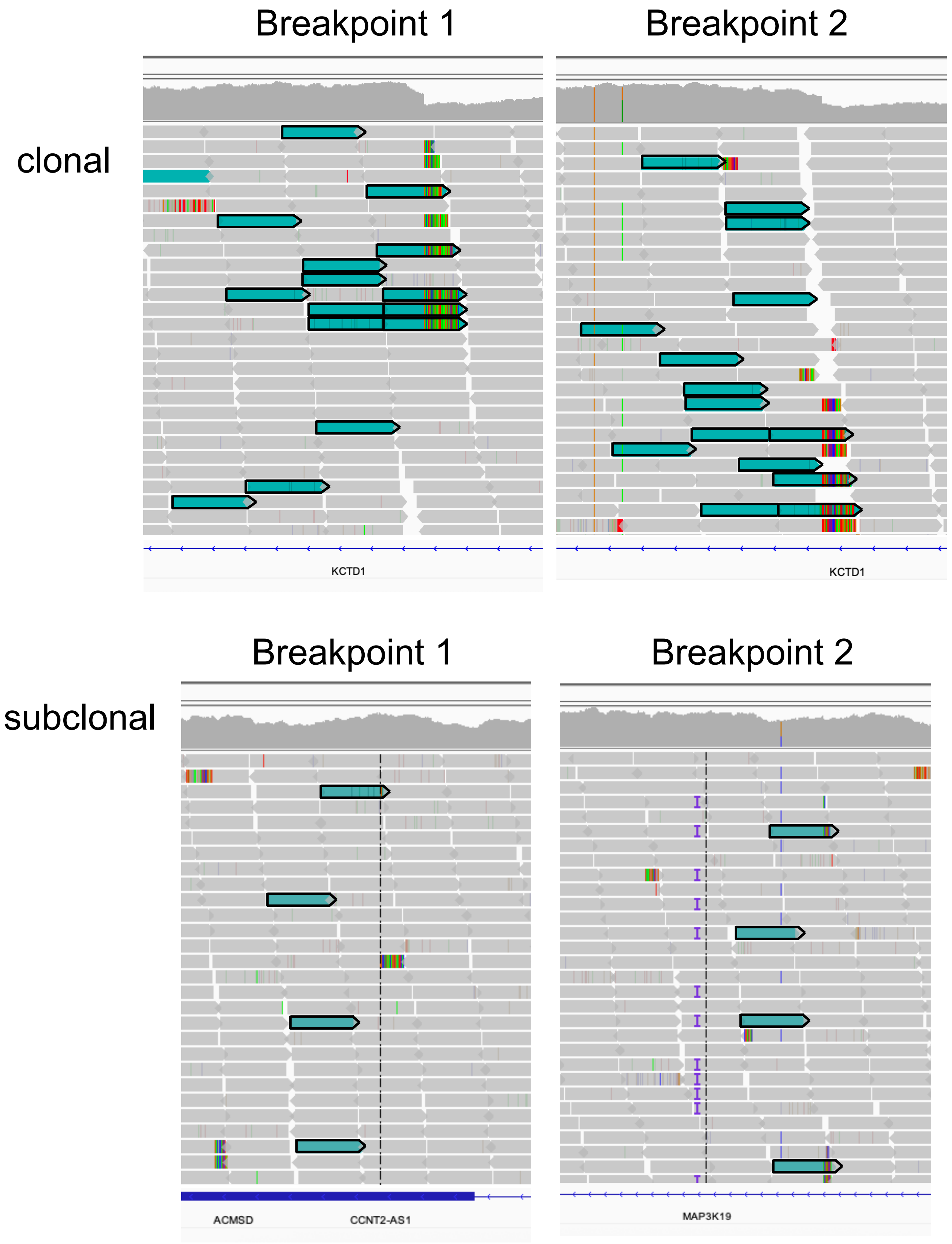

**Supplementary Figure 14 cont'd**

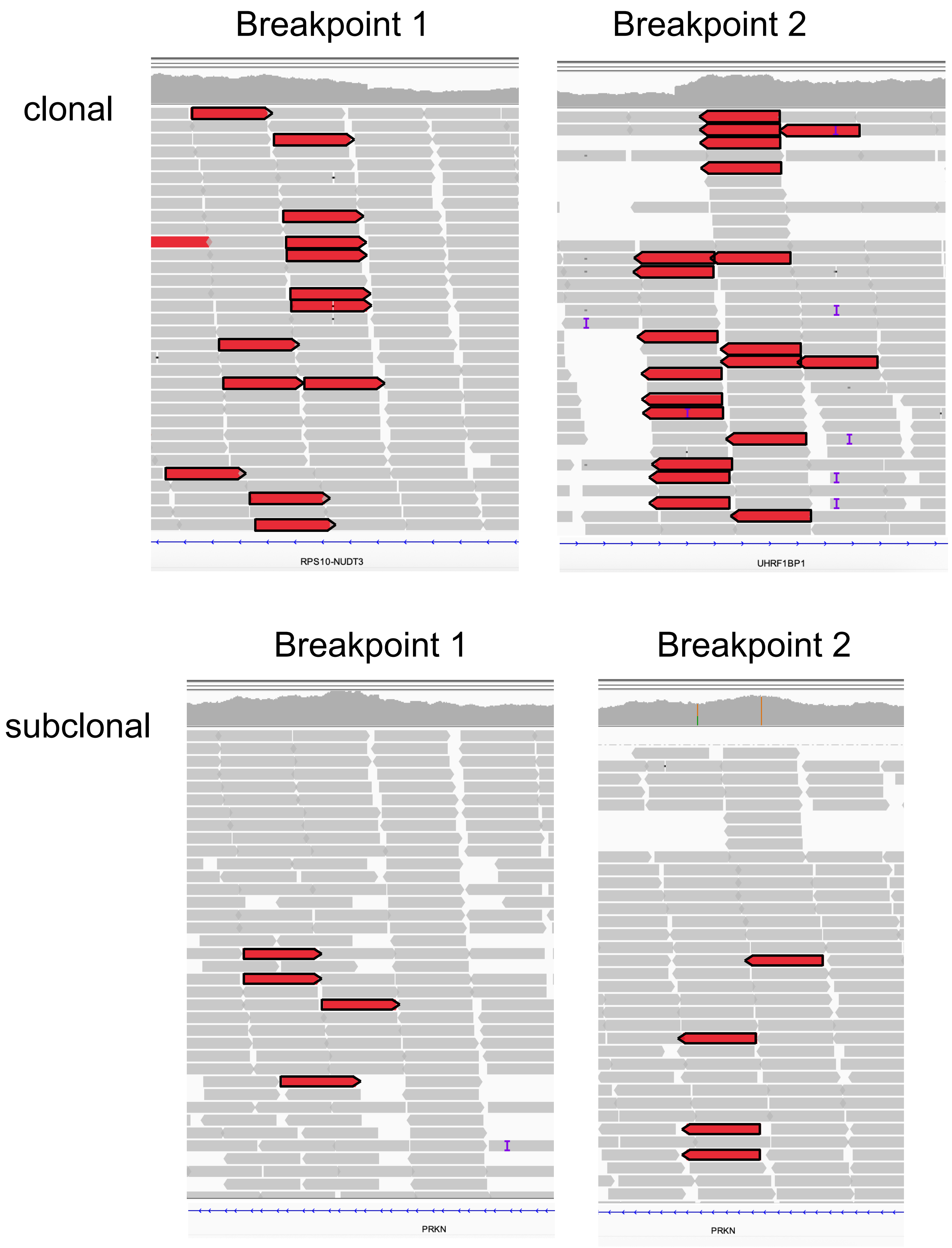

### Supplementary Figure 15

**a**

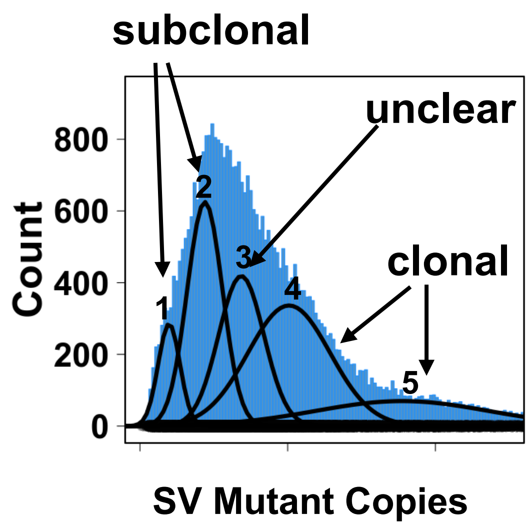


**b**

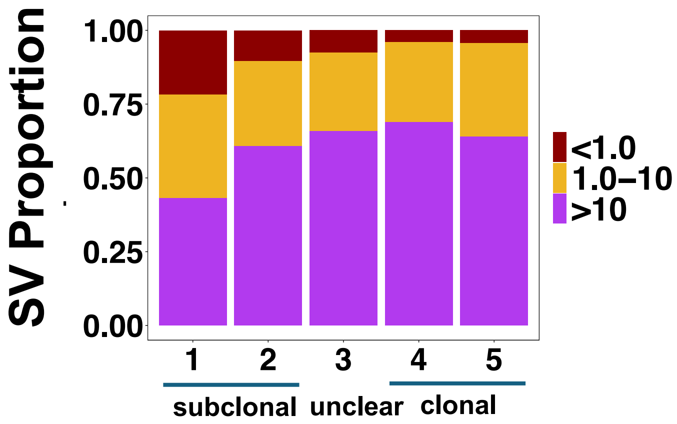

### Supplementary Figure 16

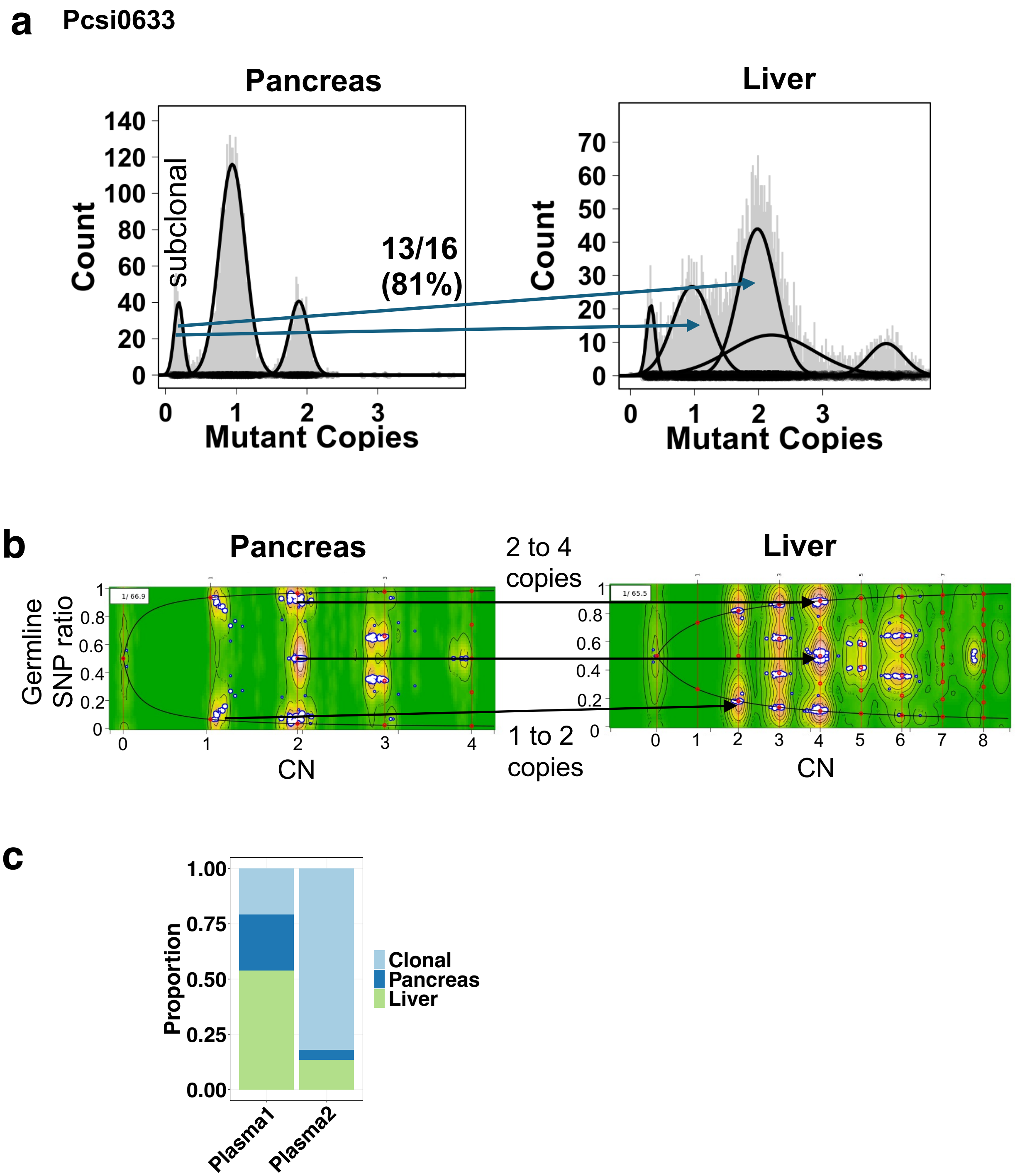

### Supplementary Figure 17

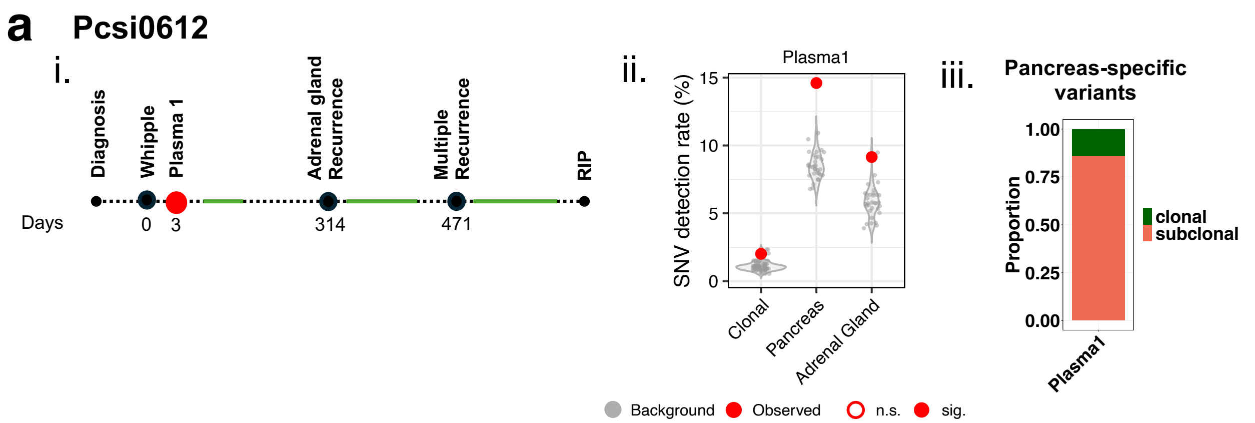


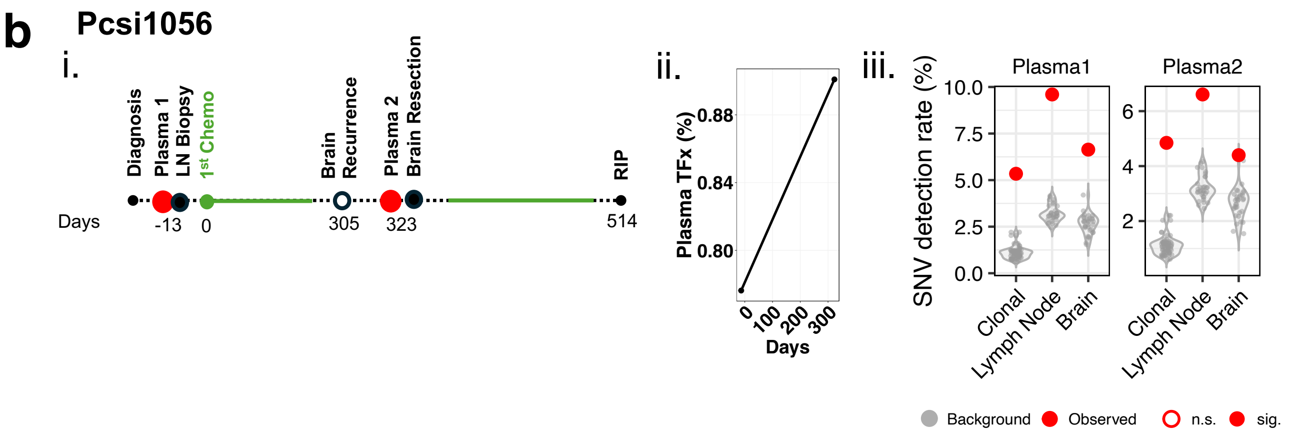


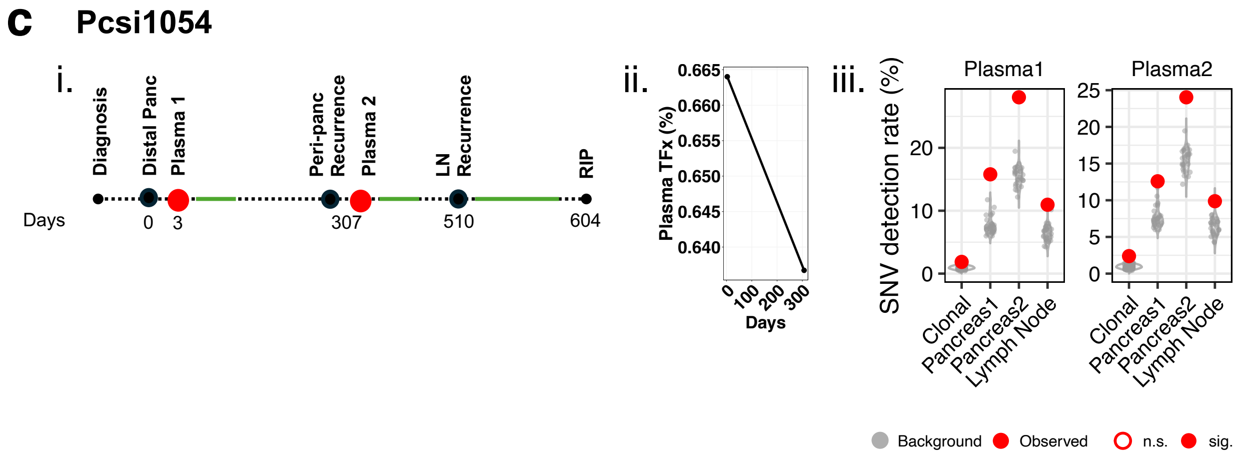

### Supplementary Figure 18

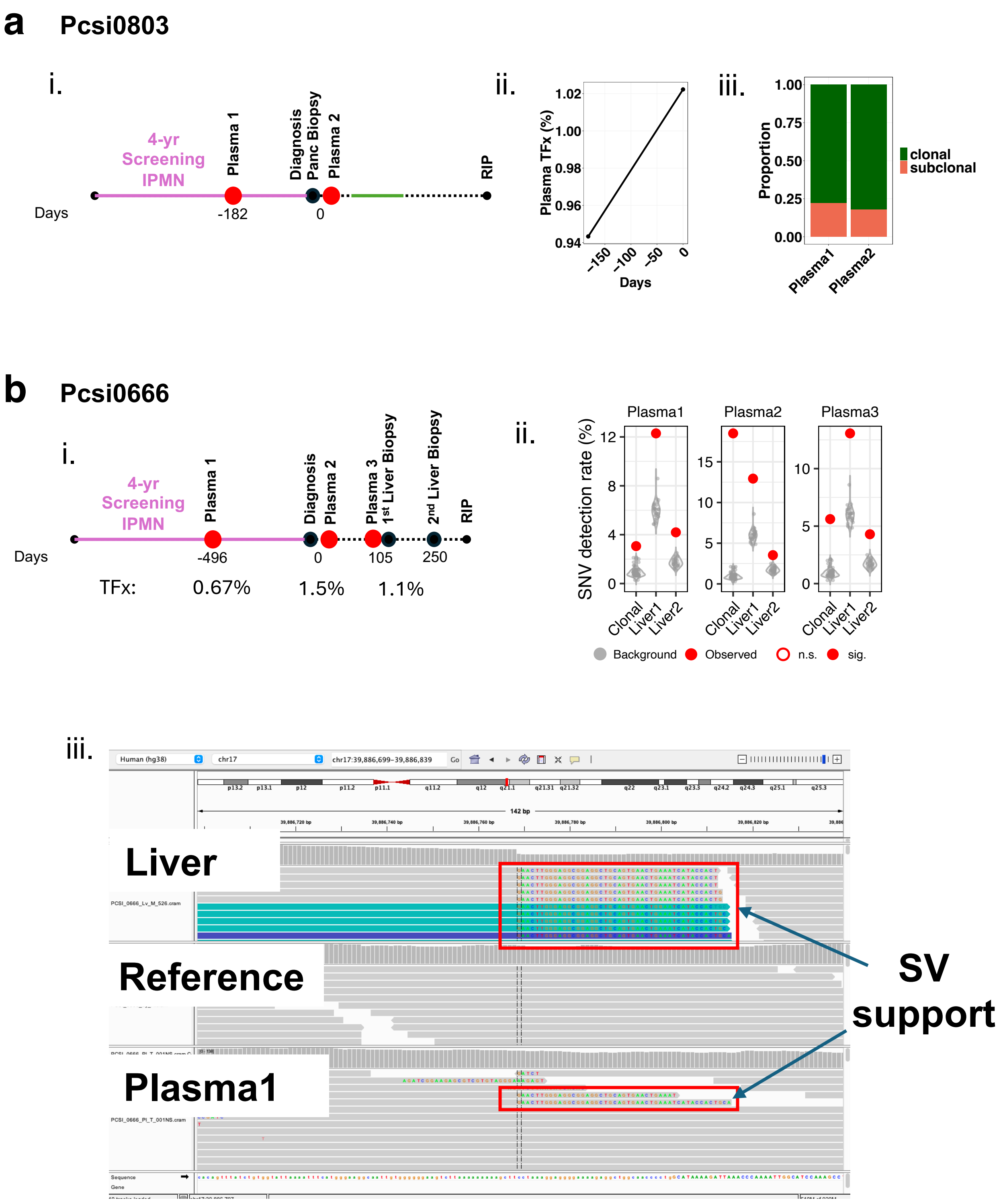

### Supplementary Figure 19

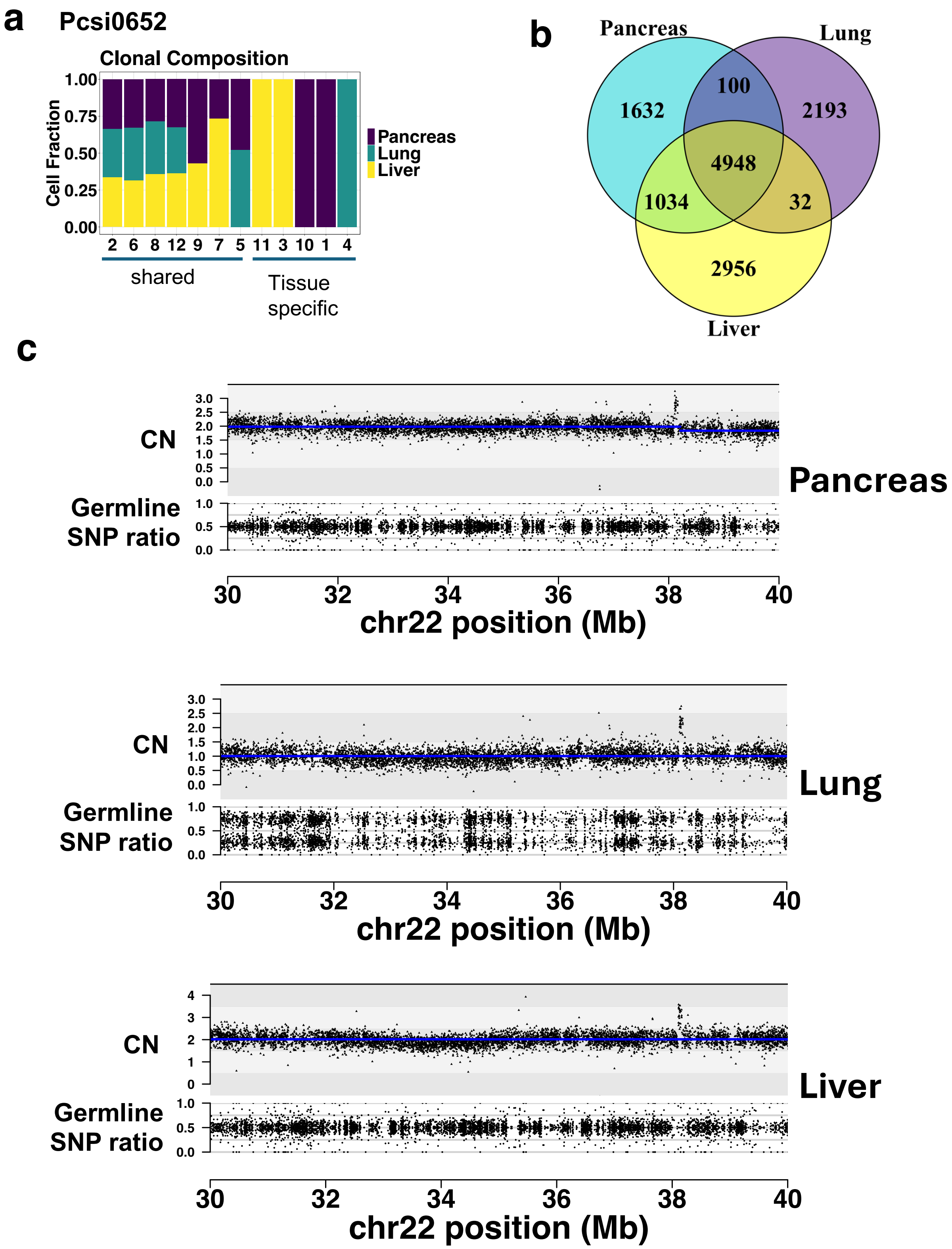

### Supplementary Figure 20

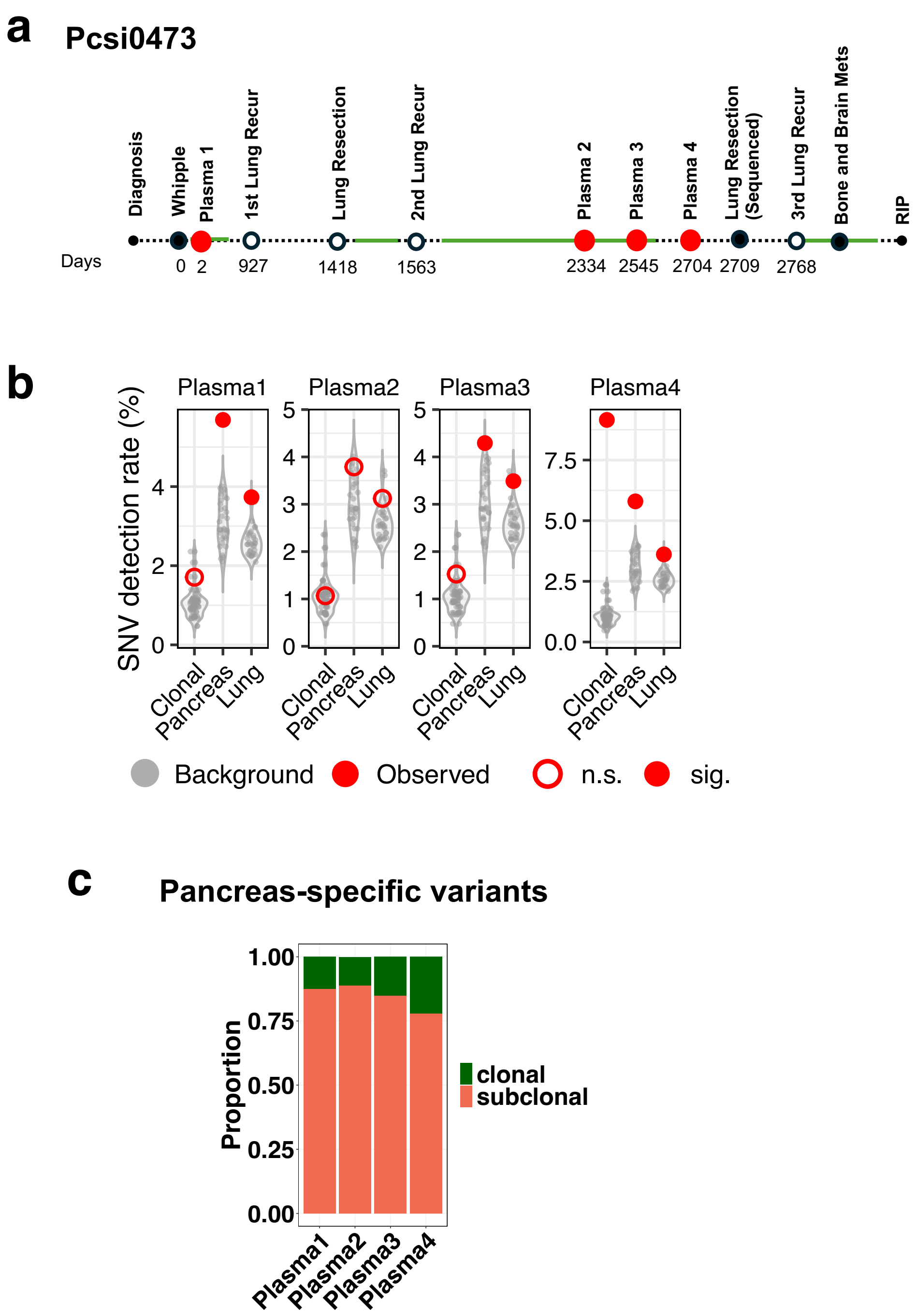

### Supplementary Figure 21

#### Pcsi0681

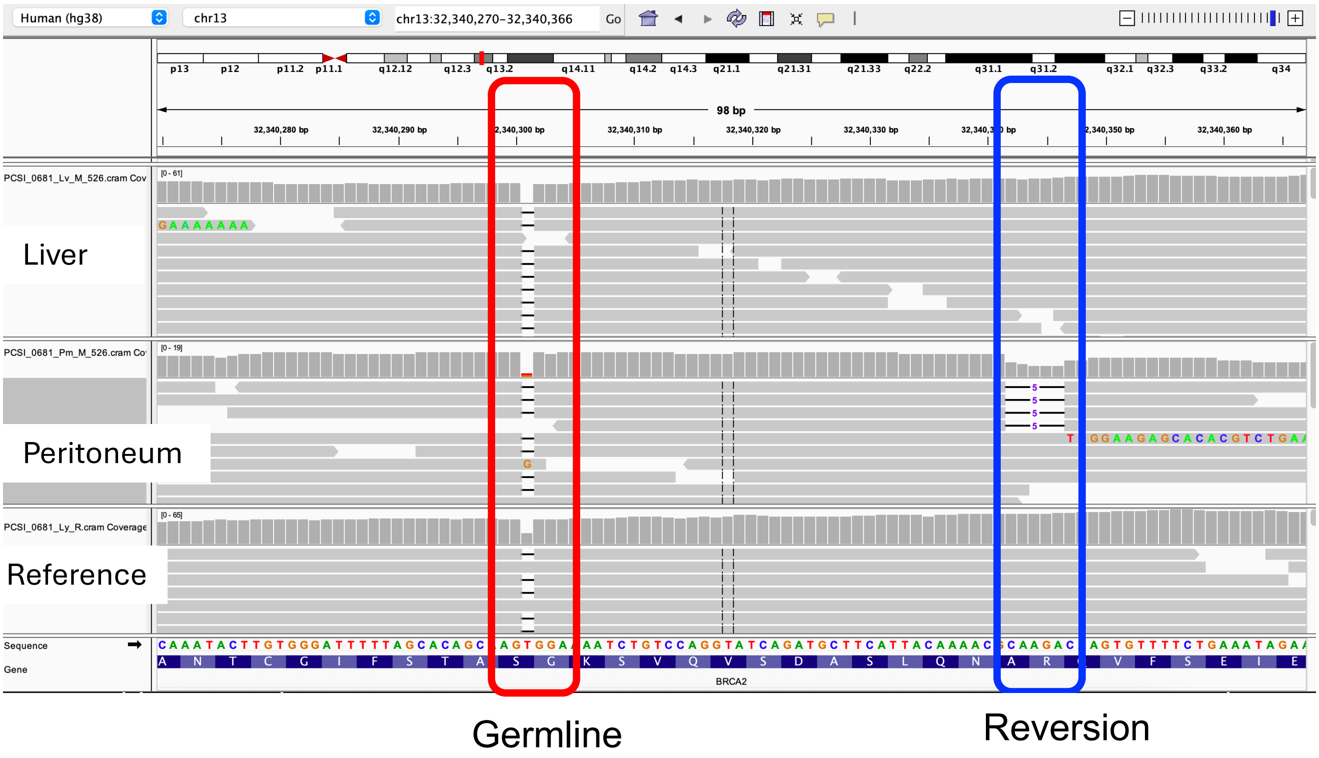
